# p107 mediated mitochondrial function controls muscle stem cell proliferative fates

**DOI:** 10.1101/2020.09.29.317693

**Authors:** Debasmita Bhattacharya, Oreoluwa Oresajo, Anthony Scimè

## Abstract

Muscle wasting diseases and aging are associated with impaired myogenic stem cell selfrenewal and a diminished number of their proliferating progenitors (MPs). Importantly, distinct metabolic states govern MP proliferation and differentiation. Central to this is the regulation between glycolysis and oxidative phosphorylation (Oxphos). However, the mechanisms that connect these energy provisioning centers to control cell behaviour remain obscure. Herein, our results reveal a mechanism by which mitochondrial-localized transcriptional co-repressor p107 governs MP proliferation. We found p107 directly interacts at the mitochondrial DNA promoter, repressing mitochondrial-encoded genes. This reduces mitochondrial ATP generation, by limiting the electron transport chain complex formation. Importantly, the amount of ATP generated by the mitochondrial function of p107 is directly associated to the cell cycle rate. Sirt1, whose activity is dependent on the cytoplasmic by-product of glycolysis, NAD^+^, directly interacts with p107 impeding its mitochondrial localization and function. The metabolic control of cell cycle, driven by differential p107 mitochondrial function, establishes a new paradigm to manipulate muscle cell proliferative fates that is likely to extend to most other dividing cell types.

## Introduction

In recent years, there has been an emerging realization that metabolism plays an active role in guiding and dictating how the cell will behave (Ghosh-Choudhary *et al*, 2020; Khacho *et al*, 2016; Porras *et al*, 2017). Central to this assertion is the interplay between metabolic states based on glucose metabolism by glycolysis in the cytoplasm with oxidative phosphorylation (Oxphos) in the mitochondria. Indeed, the cell cycle rate is dependent on the amount of total ATP generated from glycolysis and Oxphos (Martinez-Reyes *et al*, 2016; Mitra *et al*, 2009). In glycolysis, ATP is produced at a fast rate and the glycolytic pathway components might be used for the biosynthesis of nucleic acids, proteins, carbohydrates and lipids essential for cell proliferation (Folmes *et al*, 2012). On the other hand, Oxphos, which produces at least 10 times more ATP, is crucial for progression through the G1/S transition of the cell cycle (Salazar-Roa & Malumbres, 2017). Hyperactivation of Oxphos in proliferating cells is critical for their advancement, whereas its down regulation delays or blocks progression to S phase (Martinez-Reyes *et al*., 2016; Mitra *et al*., 2009; Wheaton *et al*, 2014).

Importantly, it is uncertain how glycolysis and Oxphos work together during proliferation to regulate the yield of ATP necessary for cell division. Indeed, proliferating cells customize methods to actively reduce Oxphos under conditions of higher glycolysis (Locasale & Cantley, 2011). Fundamental to understanding this relationship is the cytoplasmic NAD^+^/NADH ratio, which is a reflection of the energy generated by glycolysis. NADH is a by-product of glycolysis that might be used as a reducing reagent required for Oxphos. Whereas, NAD^+^, a coenzyme in various metabolic pathways, is the oxidized form of NADH that can be made in glycolysis from the transformation of pyruvate to lactate. High levels of cytoplasmic NAD^+^ activate the lysine sirtuin (Sirt) deacetylase family, including Sirt1, which thus acts as an energy sensor (Canto *et al*, 2015). Increased Sirt1 activity enhances mitochondrial metabolism by deacetylating histones, transcription factors and cofactors that boost mitochondrial function. In contrast, decreased NAD^+^/NADH and Sirt1 activity is associated with reduced mitochondrial function as observed in aging and metabolic diseases (Cerutti *et al*, 2014; Zhang *et al*, 2016).

In the mitochondria the capacity for ATP production is dependent on the five electron transport chain (ETC) complexes, which are made up of protein subunits derived from the nuclear and mitochondrial genomes (Martinez-Reyes & Chandel, 2020). The closed doublestranded circular mitochondrial DNA (mtDNA) encode genes located on both strands, which are referred to as the heavy (H) and light (L) strands. Their promoters are located in the large noncoding region termed the displacement loop (D-loop) (Gustafsson *et al*, 2016). The mitochondrial genome is essential to control energy generation by the ETC, as thirteen mitochondrial-encoded genes are functional components of four out of five ETC complexes. Crucially, mitochondrial genes are limiting for ATP production and inhibiting their translation can block the cell cycle in G1 (Lapuente-Brun *et al*, 2013; Martinez-Reyes *et al*., 2016; Mendelsohn *et al*, 2018; Mitra *et al*., 2009).

Mounting evidence has shown that Rb1/Rb and p107/Rbl1, members of the retinoblastoma family of transcriptional co-repressors, influence the energy status of cells and tissues (Dali-Youcef *et al*, 2007; De Sousa *et al*, 2014; Jones *et al*, 2016; Petrov *et al*, 2016; Porras *et al*., 2017; Scime *et al*, 2005; Scime *et al*, 2010; Zacksenhaus *et al*, 2017). p107 is ubiquitously expressed in proliferating cells and its over expression has been shown to block cell cycle(Wirt & Sage, 2010). However, it is also implicated in controlling stem cell and progenitor metabolic fates (De Sousa *et al*., 2014; Scime *et al*., 2005; Scime *et al*., 2010). Nonetheless, the fundamental mechanism that links p107 to metabolism and cell cycle has never been examined.

Key to understanding this relationship are the muscle resident myogenic stem cells (satellite cells) SCs, which are characterized by dynamic metabolic reprogramming during different stages of the differentiation process (Bhattacharya & Scime, 2020). They use predominately Oxphos in quiescence, but their committed progeny, the myogenic progenitor (MP) cells up-regulate glycolysis to support proliferation (Bhattacharya & Scime, 2020; Ryall *et al*, 2015). In this study, MPs were used to uncover a novel control mechanism of energy generation that showcases the interplay between glycolysis and Oxphos in regulating cell proliferation. Intriguingly, we found that p107 accomplishes this by controlling ATP generation capacity in the mitochondria through a Sirt1 dependent mechanism. Thus, our findings highlight a crucial role for p107 linking the regulation of mitochondrial Oxphos to proliferation dynamics in MPs that is likely to extend to other cell types.

## Results

### p107 is localized in the mitochondria of myogenic progenitor (MP) cells

By Western blotting cytoplasmic and nuclear cellular fractions, p107 was found in the cytoplasm of actively proliferating MP cell line c2c12 (c2MPs), as is the case in other cell types (Wirt & Sage, 2010) (**Fig. 1A**). Considering that emergent findings show that p107 can influence the metabolic state of progenitors (Porras *et al*., 2017), we assessed its potential metabolic role in the cytoplasm. As mitochondria are crucial in controlling metabolism, we assessed for the presence of p107 within this organelle. Intriguingly, Western blot analysis demonstrated that p107 was in the mitochondria fraction of the cytoplasm during proliferation, but almost absent when the cells were contact inhibited in confluent growth arrested cultures (**Fig. 1B**). To further assess for the presence of p107, we isolated various mitochondrial fractions utilizing hypotonic osmotic shock by varying sucrose concentrations (Lu *et al*, 2009). Mitochondrial fractionation showed that p107 was not located at the outer membrane nor within the soluble inner membrane fraction whose soluble proteins were relinquished by decreasing the buffer molarity (**Fig. 1C**). However, it was localized within the matrix where the protein constituents are unavailable to trypsin for digestion compared to the soluble inner membrane proteins such as Cox4, as well as p107 from the whole cell lysate control (**Fig. 1C**).

**Fig 1.**
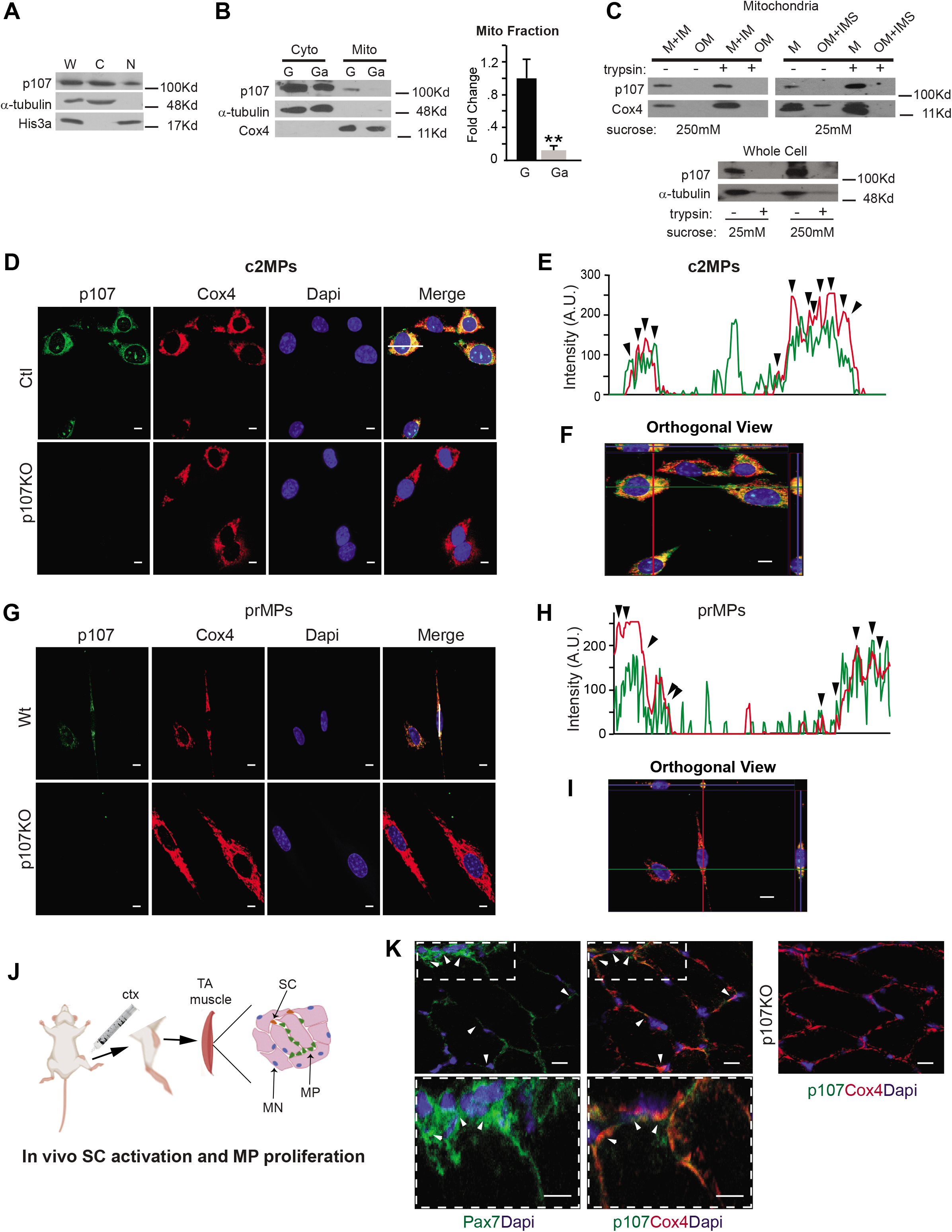
p107 is localized in the mitochondria of proliferating myogenic progenitors. **(A)** Representative Western blot of whole cell (W), cytoplasmic (C) and nuclear (N) fractions for p107, α-tubulin (cytoplasm loading control) and His3a (nucleus loading control) during proliferation. **(B)** Representative Western blot and graphical representation of cytoplasmic (Cyto) and mitochondrial (Mito) c2MP fractions for p107, α-tubulin and Cox4 (mitochondria loading control) during proliferation (G) and growth arrest (Ga), n=3, asterisks denote significance, \*\**p*<0.01, Student T-test. **(C)** Representative Western blot of c2MP whole cell and mitochondrial fractions including outer membrane (OM), inner membrane (IM), soluble inner membrane (IMS) and matrix (M) that were isolated in 250mM or 25mM sucrose buffer, treated and untreated with trypsin. Confocal immunofluorescence microscopy for p107, Cox4, Dapi and Merge of proliferating **(D)** Control (Ctl) and genetically deleted p107 (p107KO) c2MPs and **(G)** wild type (Wt) and p107KO primary (pr) MPs (scale bar 10μm). **(E)** and **(H)** A line was drawn through a representative cell to indicate relative intensity of RGB signals with the arrowheads pointing to areas of concurrent intensities **(F)** and **(I)** an orthogonal projection was generated by a Z-stack (100 nm interval) image set using the ZEN program (Zeiss) in the XY, XZ, and YZ planes. **(J)** Schematic for inducing in vivo satellite cell (SC) activation and MP proliferation in the tibialis anterior (TA) muscle with cardiotoxin (ctx) injury (MN is myonuclei). **(K)** Confocal immunofluorescence microscopic merged image of wild type (Wt) TA muscle section 2 days post injury for Pax7 and Dapi and p107, Cox4 and Dapi and for p107KO TA muscle section for p107, Cox4and Dapi (scale bar 20μm). Arrows denote Pax7 ^+^p107^+^Cox4^+^ cells (scale bar 20μm).

These results were corroborated by confocal microscopy and subsequent analysis of generated z-stacks that showed p107 and the mitochondrial protein Cox4 co-localize in c2MPs **(Fig. 1D)**. The specificity of immunocytochemistry was confirmed by immunofluorescence of p107 in p107 “knockout” c2MPs (p107KO c2MPs) generated by Crispr/Cas9 **(Fig. 1D & Suppl. Fig. 1A)**. Moreover, p107 and Cox4 fluorescence intensity peaks were matched on a line scanned RGB profile **(Fig. 1E)** and orthogonal projection showed co-localization in the XY, XZ, and YZ planes (**Fig 1F)**. The same assays were used to confirm these findings in proliferating primary wild type (Wt) MPs (prMPs) and p107 genetically deleted prMPs (p107KO prMPs) isolated from Wt and p107KO mouse skeletal muscles, respectively **(Fig. 1G, 1H, 1I)**. In vivo, in a model of regenerating skeletal muscle caused by cardiotoxin injury, we also found p107 co-localized with Cox 4 in proliferating MPs, which express the myogenic stem cell marker Pax7 (**Fig. 1J, 1K & Suppl. Fig. 2).** Together, these data showcase that p107 localizes in the mitochondria of MPs from primary and muscle cell lines, suggesting that it might have an important mitochondrial function in the actively dividing cells.

### p107 interacts at the mtDNA

To find the functional consequence of p107 mitochondrial matrix localization we assessed if it interacts at the mtDNA to repress mitochondrial gene expression similar to its role as a co-transcriptional repressor on nuclear promoters (Wirt & Sage, 2010). We evaluated this potential by performing quantitative chromatin immunoprecipitation (qChIP) analysis on the D-loop regulatory region of isolated mitochondria from c2MPs. qChIP revealed that p107 interacted at the mtDNA of growing c2MPs (**Fig. 2A**). It was negligible in growth arrested cells when p107 levels are deficient in the mitochondria and not detected in p107KO c2MPs (**Fig. 2A**).

**Fig 2.**
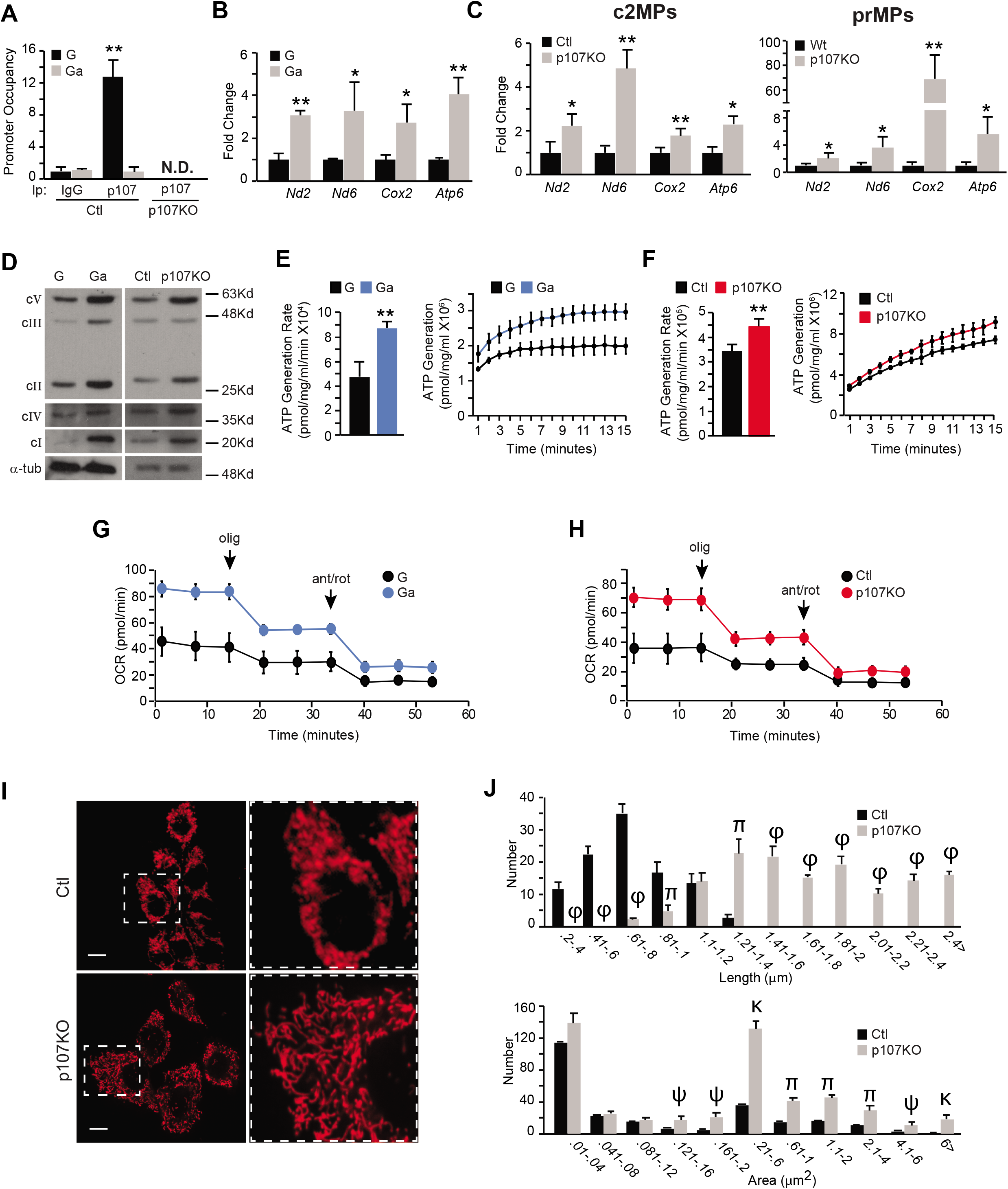
p107 down regulates mitochondrial ATP generation capacity. **(A)** Graphical representation of relative p107 and IgG mitochondrial DNA promoter occupancy by qChIP analysis during proliferation (G) and growth arrest (Ga) in control (Ctl) and p107KO c2MPs, n=3; \*\**p*<0.01; two-way Anova with post hoc Tukey. Not detected (N.D.). Gene expression analysis by qPCR of mitochondrial encoded genes *Nd2, Nd6, Cox2* and *Atp6* for **(B)** c2MPs during G and Ga, n=3 and **(C)** Ctl and p107KO c2MPs n=4, and Wt and p107KO prMPs during proliferation, n=4; \**p*<0.05; \*\**p*<0.01; Student T-test. **(D)** Representative Western blot of G compared to Ga, and Ctl compared to p107KO c2MPs for each electron transport chain complex (c) subunits, cI (NDUFB8), cII (SDHB), cIII (UQCRC2), cIV (MTCO) and cV (ATP5A). **(E**) ATP generation rate and capacity over time for isolated mitochondria from G and Ga c2MPs n=4 and for **(F)** Ctl and p107KO c2MPs, n=4. For rate, \*\**p*<0.01, Student T-test. For capacity significance statistics see **Suppl. Table 1A and 1B**. Live cell metabolic analysis by Seahorse for oxygen consumption rate (OCR) with addition of oligomycin (olig) and antimycin A (ant), rotenone (rot) for **(G)** G and Ga c2MPs n=6-8 and **(H)** Ctl and p107KO c2MPs, n=4. **(I)** Representative confocal microscopy live cell image of stained mitochondria (MitoView Red) for Ctl and p107KO c2MPs. Graphical representation of mitochondria number per grouped **(J)** lengths and areas using image J quantification; n=3; significance denoted by ψ=p<0.05, π=p<0.01, κ=p<0.001, ϕ=p<0.0001; Student T-test.

### p107 represses mtDNA encoded gene expression to regulate Oxphos capacity

We next appraised if p107 mtDNA promoter occupancy might repress mitochondrial encoded gene expression. For this we assessed 4 of 13 mitochondrial genes that are subunits of 4 of 5 ETC complexes. qPCR analysis showed that during proliferation when p107 is abundant in the mitochondria, there was significantly less expression of the mitochondrial encoded genes as compared to growth arrested cells (**Fig. 2B**). The importance of p107 to mitochondrial gene expression was confirmed with p107KO c2MPs and prMPs, which both exhibited significantly increased mitochondrial encoded gene expression in the genetically deleted cells compared to their controls (**Fig. 2C**). The increased mitochondrial gene expression and Oxphos in growth arrested and p107KO cells corresponded to augmented ETC complex formation revealed by Western blot analysis of the relative protein levels of the 5 Oxphos complexes **(Fig. 2D).**

As mitochondrial gene expression is limiting for ETC complex formation, we used two methods to gauge if p107 promoter occupancy was associated with the capacity of mitochondria to generate ATP. First, using a luminescence assay, ATP generation potential capacity and rate were determined from isolated mitochondria, which were normalized to their protein content to exclude the potential contribution of mitochondria number. We found that the potential rate and capacity of ATP formation from isolated mitochondria of growth arrested cells and p107KO c2MPs were significantly higher than proliferating and control c2MPs, respectively **(Fig. 2E, 2F & Suppl. Table 1A)**, corresponding to the increase in mitochondrial gene expression profile and ETC complex formation **(Fig. 2B, 2C & 2D)**.

Second, we confirmed these results with live cell metabolic analysis using Seahorse, which showed that both growth arrested cells and p107KO c2MPs had significantly enhanced production of mitochondrial ATP generated compared to the proliferating and control c2MPs, respectively **(Fig. 2G, 2H & Suppl. Fig. 3)**. As Oxphos is known to increase mitochondrial fusion causing elongation, we assessed a potential effect on mitochondrial morphology (Mishra *et al*, 2014; Mishra & Chan, 2016). Interestingly, we found that p107KO c2MPs had an elongated mitochondrial network **(Fig. 2I)** made up of mitochondria with significantly increased length and area **(Fig. 2J)**. Together these data suggest that p107 has repressor activity when bound to the mtDNA promoter reducing the capacity to produce ETC complex subunits, which influences the mitochondria potential for ATP generation.

### Glycolytic flux controls p107 mitochondrial function

The reliance of glycolysis versus Oxphos during the cell cycle can be driven by nutrient accessibility. Thus, we evaluated how nutrient availability influences p107 compartmentalization during proliferation. We assessed the effect of glucose metabolism on p107 mitochondrial localization by growing c2MPs in stripped media with glucose as the sole nutrient. Western blot analysis of isolated mitochondria revealed that p107 translocation into mitochondria was glucose concentration dependent (**Fig. 3A)**. Its presence was negligible in mitochondria when cells were grown in 5.5mM glucose, but there was a significant increase of p107 localization in mitochondria at a higher glucose concentration (25mM). The increased level of mitochondrial p107 in c2MPs grown in higher glucose concentration corresponded to significantly increased mtDNA binding at the D-loop (**Fig. 3B**). This was associated with a significant decrease in mitochondrial encoded gene expression in both c2MPs and prMPs grown in high glucose concentration (**Fig. 3C & Suppl. Fig. 4**). However, increasing the glucose concentration in the stripped media had no effect on mitochondrial gene expression in the p107KO c2MPs and p107KO prMPs compared to their controls (**Fig. 3C & Suppl. Fig. 4**).

**Fig 3.**
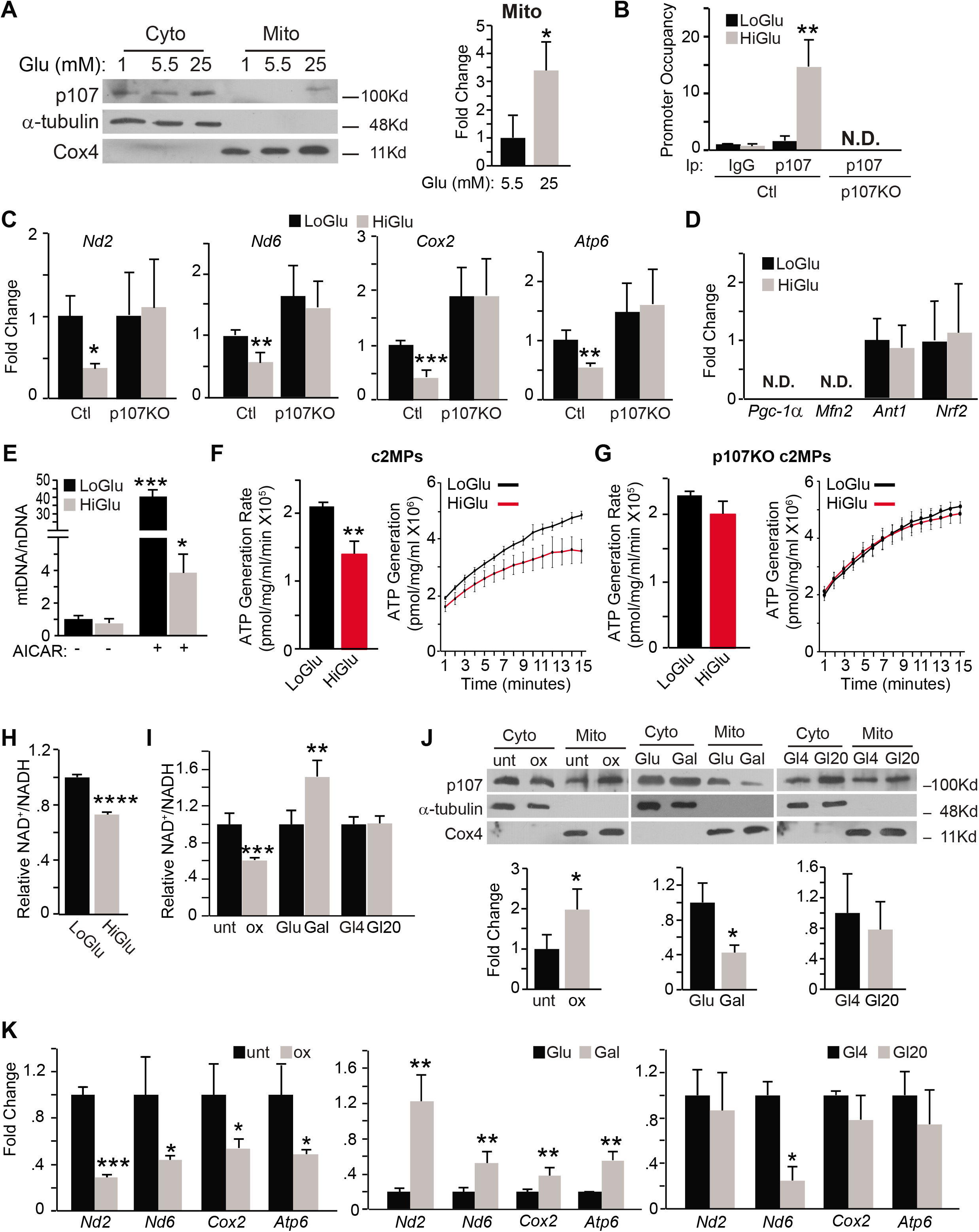
p107 discerns the NAD^+^/NADH ratio to regulate Oxphos. **(A)** Representative Western blot and graphical representation of cytoplasmic (Cyto) and mitochondrial (Mito) fractions of c2MPs grown in stripped media (SM) containing only 1.0mM, 5.5mM or 25mM glucose (Glu), n=3, \**p*<0.05; Student T-test. **(B)** Graphical representation of relative p107 and IgG mitochondrial DNA promoter occupancy by qChIP analysis for control (Ctl) and p107KO c2MPs grown in SM containing only 5.5mM (LoGlu) or 25mM (HiGlu) glucose, n=3, \*\**p*<0.01; two-way Anova with post hoc Tukey. **(C)** qPCR of *Nd2, Nd6, Cox2* and *Atp6* for Ctl and p107KO c2MPs grown in SM containing LoGlu or HiGlu, n=4, \**p*<0.05, \*\**p*<0.01, \*\*\**p*<0.001; two-way Anova with post hoc Tukey. **(D)** qPCR of *Pgc-1α, Mfn2, Ant1* and *Nrf2* for c2MPs grown in SM containing LoGlu or HiGlu, n=3 (N.D. is not detected). **(E)** Mitochondrial DNA (mtDNA) to nuclear DNA (nDNA) ratio for c2MPs grown in SM containing LoGlu or HiGlu in the absence or presence of 5-aminoimidazole-4-carboxamide ribonucleotide (AICAR), n= 3. \**p*<0.05, \*\*\**p*<0.001; two-way Anova with post hoc Tukey. Isolated mitochondria ATP generation rate and capacity over time for **(F)** c2MPs (n =4) and **(G)** p107KO c2MPs (n =4), grown in SM containing LoGlu or HiGlu. For rate, \*\**p*<0.01; Student T-test. For capacity significance statistics see Suppl. Table 1C. NAD^+^/NADH ratio for **(H)** c2MPs grown in SM containing LoGlu or HiGlu. n=3 and **(I)** c2MPs untreated (Unt) or treated with 2.5mM oxamate (ox), grown in 25mM glucose (Glu) or 10mM galactose (Gal) or with varying glutamine concentrations of 4mM (Gl4) or 20mM (Gl20), n=4, \*\**p*<0.01, \*\*\**p*<0.001, \*\*\*\**p*<0.0001; Student T-test. **(J)** Representative Western blots and graphical representation of mitochondrial (Mito) fractions of cells in (I), n=3, \**p*<0.05; Student T-test. **(K)** qPCR analysis of cells in (I) for *Nd2, Nd6, Cox2* and *Atp6*. n=3, \**p*<0.05, \*\**p*<0.01, \*\*\**p*<0.001; Student T-test.

Importantly, our results also verified that changes in mitochondrial gene expression in MPs grown in varying glucose concentrations is not influenced by differences in mitochondrial biogenesis. By using qPCR, the gene expression of key markers of mitochondrial biogenesis *Pgc-1α* and *Mfn2* were not detected and there were no differences for expression of *Ant1* and *Nrf2* in c2MPs grown in 5.5mM compared to 25mM glucose (**Fig. 3D**). Moreover, the mtDNA to nuclear DNA ratio remained unchanged when grown in the different glucose concentrations, in contrast to MPs treated with the mitochondrial biogenesis activator AICAR (**Fig. 3E**). Together, these data show that mitochondrial gene expression might be mediated by glycolytic flux induced p107 mitochondrial gene repression and not merely a consequence of differences in mitochondrial biogenesis.

We next investigated if the reduction of mitochondrial gene expression with increased glucose availability influenced the potential mitochondrial ATP synthesis. In isolated mitochondria, we determined that there was a significant decrease in the potential rate and capacity of ATP generation when cells were grown in 25mM compared to 5.5mM glucose (**Fig. 3F & Suppl. Table 1C)**. This corresponds to when p107 is in the mitochondria compared to low glucose when it is barely present. However, in the p107KO c2MPs, where mitochondrial gene expression was not influenced by glucose (**Fig. 3C**), the potential mitochondrial ATP rate and generation capacity also was unaffected by glucose **(Fig. 3G**). These data suggest that p107 is controlled by glucose metabolism to influence the potential mitochondrial energy production through regulation of the mitochondrial gene expression.

### NAD^+^/NADH regulates p107 mitochondrial function

As glucose metabolism affects the NAD^+^/NADH redox balance, we determined if this might impact p107 mitochondrial localization and function. We found that c2MPs grown in 5.5mM compared to 25mM glucose in stripped media had significantly increased cytoplasmic NAD^+^/NADH ratio (**Fig. 3H**). This suggests that the cytoplasmic NAD^+^/NADH ratio might influence mitochondrial gene expression and ATP generation capacity via a p107 dependent mechanism.

We manipulated the cytoplasmic NAD^+^/NADH energy flux in a glucose concentration independent manner to find the importance of the redox potential to p107 function. We added oxamate (ox) or dichloroacetic acid (DCA) to decrease the NAD^+^/NADH ratio or grew cells in the presence of galactose instead of glucose, which increases NAD^+^/NADH ratio (**Fig. 3I and Suppl. Fig 5A**). We also grew cells in varying amounts of glutamine in stripped media, which had no effect on the cytoplasmic NAD^+^/NADH ratio (**Fig. 3I)**. As anticipated, Western blot analysis of c2MPs treated with ox and DCA showed significantly increased p107 levels in the mitochondria (**Fig. 3J and Suppl. Fig. 5B**). On the other hand, galactose treated cells had significantly less p107 in the mitochondria compared to untreated controls and cells treated with glutamine showed no difference in the level of p107 mitochondrial localization (**Fig. 3J**). These results suggest that p107 mitochondrial localization is based on the cytoplasmic NAD^+^/NADH ratio. Mitochondrial gene expression levels with the different treatments corresponded with p107 mitochondrial localization established by the cytoplasmic NAD^+^/NADH ratio (**Fig. 3K and Suppl. Fig. 5C**). Indeed, significantly higher mitochondrial gene expression levels were observed with higher cytoplasmic NAD^+^/NADH when cells were grown with galactose compared to glucose, whereas significantly lower mitochondrial gene expression was evident when cells were treated with ox and DCA that caused decreased cytoplasmic NAD^+^/NADH (**Fig. 3K and Suppl. Fig. 5C**). No significant gene expression changes were present when cells were grown solely with varying amounts of glutamine, except *Nd6*, which is the only mitochondrial gene expressed from the L-strand of mtDNA (**Fig. 3K**). Together, these results show that p107 acts indirectly as an energy sensor of the cytoplasmic NAD^+^/NADH ratio that might influence the potential ATP produced from the mitochondria, as a consequence of regulating mitochondrial gene expression.

### Sirt1 directly regulates p107 mitochondrial function

As the NAD^+^ dependent Sirt1 deacetylase is an energy sensor of the NAD^+^/NADH ratio, we evaluated if it potentially regulates p107. Importantly, reciprocal Immunoprecipitation/Western analysis on c2MP lysates for endogenous p107 and Sirt1 showed that they directly interacted (**Fig. 4A)**. No interactions were apparent in p107KO and Sirt1genetically deleted (Sirt1KO) c2MPs obtained by Crispr/Cas9 (**Fig. 4A and Suppl Fig. 1B**).

**Fig 4.**
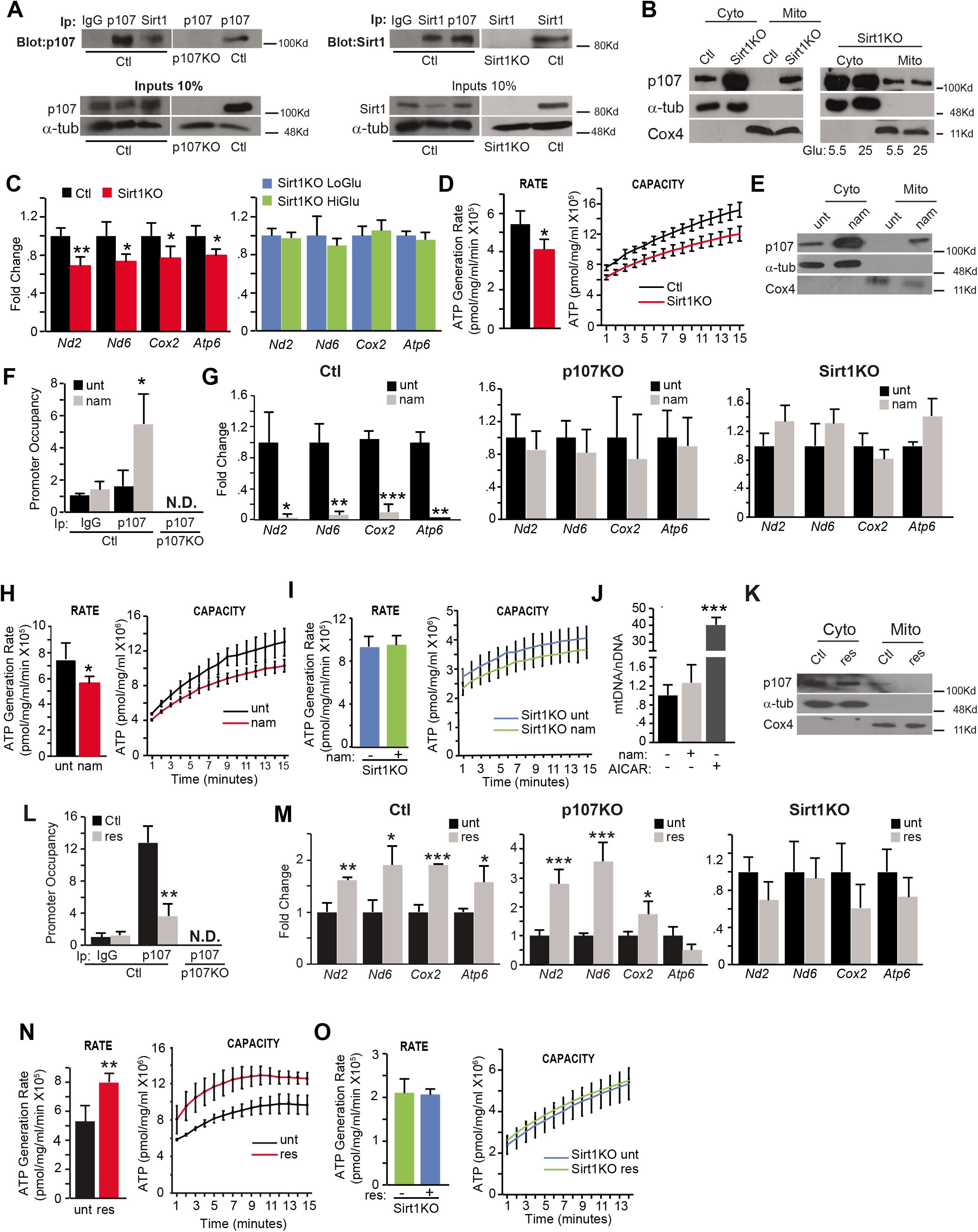
p107 is regulated by Sirt1. **(A)** Immunoprecipitation/Western blots for p107 and Sirt1 in control (Ctl) and p107KO c2MPs. (**B)** Representative Western blot of cytoplasmic (Cyto) and mitochondrial (Mito) fractions of Ctl and Sirt1KO c2MPs grown in stripped media with 5.5mM and 25mM glucose. **(C)** qPCR of *Nd2, Nd6, Cox2* and *Atp6* for Ctl and Sirt1KO c2MPs grown in stripped media with 5.5mM (LoGlu) and 25mM (HiGlu) glucose, n=4; \**p*<0.05, \*\**p*<0.01 **(D)** Isolated mitochondrial ATP generation rate and capacity over time for Ctl and Sirt1KO c2MPs grown in 5.5mM glucose. For rate n=4, \**p*<0.05. For capacity significance statistics see Suppl. Table 1D **(E)** Representative Western blot of Cyto and Mito fractions of c2MPs untreated (Ctl) or treated with (10mM) Sirt1 inhibitor nicotinamide (nam) or **(K)** a concentration (10μM) of resveratrol (res) that activates Sirt1. Graphical representation of relative p107 and IgG mitochondrial DNA promoter occupancy by qChIP analysis in Ctl and p107KO c2MPs untreated (unt) or treated with **(F)** nam (n=3) and **(L)** a concentration of res (10μM) that activates Sirt1 (n=3), \**p*<0.05, \*\**p*<0.01; two-way Anova with post hoc Tukey. qPCR of *Nd2, Nd6, Cox2* and *Atp6* for Ctl, p107KO and Sirt1KO c2MPs unt and treated for **(G)** nam (n=4) and **(M)** 10μM res, (n=3-4); \**p*<0.05, \*\**p*<0.01, \*\*\**p*<0.001; Student T-test. Isolated mitochondria ATP generation rate and capacity over time for **(H)** Ctl and **(I)** Sirt1KO c2MPs treated and untreated with nam, n=4 or **(N)** Ctl and **(O)** Sirt1KO c2MPs treated or untreated with 10μMres, n=4. For rate \**p*<0.05, \*\**p*<0.01; Student T-test. For capacity significance statistics see Suppl. Table 1E and 1F. **(J)** Mitochondria DNA (mtDNA) to nuclear DNA (nDNA) ratio for c2MPs with and without AICAR and nam, n=3-4, \*\*\**p*<0.001; Student T-test.

We next assessed if Sirt1 activity affected p107 mitochondrial function. Unlike c2MPs grown in 5.5mM glucose that exhibit relocation of p107 from the mitochondria, Sirt1KO cells did not alter p107 mitochondrial localization **(Fig. 4B & 3A).** This resulted in significantly decreased mtDNA encoded gene expression for Sirt1KO cells grown in 5.5mM glucose compared to controls and no differences between Sirt1KO cells grown in 5.5mM or 25mM glucose **(Fig. 4C)**. The decreased mitochondrial gene expression in Sirt1KO cells grown in 5.5mM glucose corresponded with a reduced mitochondrial ATP generation rate and capacity compared to control cells (**Fig. 4D & Suppl. Table 1D)**. There were no differences in ATP generation rate or capacity between Sirt1KO cells grown in 5.5mM or 25mM glucose **(Suppl. Fig. 6**).

We next determined if Sirt1 activity is necessary for p107 mitochondrial function. Inhibition of Sirt1 activity by nicotinamide (nam) increased p107 mitochondrial localization (**Fig. 4E**) that was concomitant with increased mtDNA promoter interaction (**Fig. 4F**). This was associated with decreased mitochondrial gene expression, whereas nam treatment of p107KO and Sirt1KO c2MPs had no effect on the mitochondrial gene expression (**Fig. 4G**). This suggests that Sirt1 control of mitochondrial gene expression is dependent on p107. As expected, both the potential ATP generation rate and capacity of isolated mitochondria were reduced with Sirt1 attenuated activity (**Fig. 4H & Suppl. Table 1E**). On the contrary, Sirt1KO c2MPs showed no difference in ATP generation capacity and rate, and no changes in mitochondrial gene expression profile (**Fig. 4I**). Moreover, the reduction of Oxphos capacity by attenuated Sirt1 activity was not due to an apparent decrease in mitochondrial biogenesis, as nam treatment did not influence the mtDNA/nDNA ratio (**Fig. 4J)**.

To confirm these results, we also used resveratrol (res) at low and high doses that indirectly activate and inhibit Sirt1 activity, respectively. c2MPs grown in a low concentration of res (10μM), had the opposite effect to nam for p107 localization and function. Treatment of c2MPs with this concentration of res decreased p107 within the mitochondria (**Fig. 4K)**. The activation of Sirt1 also reduced p107 mtDNA promoter interaction and enhanced the mitochondrial gene expression (**Fig. 4L & 4M**), which corresponded to an increased ATP synthesis rate and capacity of isolated mitochondria (**Fig. 4N & Suppl. Table 1F**). No differences in ATP generation rate and capacity were observed in Sirt1KO c2MPs, which was anticipated with a non-significant consequence on mitochondrial gene expression (**Fig. 4M & 4O**). When Sirt1 activity was repressed by high doses of res, p107 was localized in the mitochondria and gene expression along with ATP generation rate and capacity were significantly decreased (**Suppl. Fig. 7 & Suppl. Table 1G**).

We further assessed the role of Sirt1 activity on p107 localization by overexpressing c2MPs with Ha tagged p107 in the presence of full length Sirt1 (Sirt1fl) or dominant negative Sirt1(Sirt1dn). Similar to Sirt1KO cells, Sirt1dn had no effect on preventing p107 mitochondria localization when grown in 5.5mM glucose as opposed to Sirt1fl **(Compare Fig. 4B & Suppl. Fig. 8).** As expected, the Sirt1fl overexpression did not alter p107 localization in cells grown in 25 mM glucose (**Suppl. Fig. 9).** Also, there was significantly less mitochondrial encoded gene expression in Sirt1dn cells grown in 5.5mM glucose compared to Sirt1fl **(Compare Fig 4C & Suppl Fig. 10)**. Together, these results suggest that p107 is influenced directly by Sirt1 activity to localise within the cytoplasm, de-repressing mitochondrial gene expression and hence increasing ATP generation capacity.

### p107 directs cell cycle rate through management of Oxphos generation

We next considered how this metabolic role for p107 might influence MP behaviour in vivo by assessing skeletal muscle regeneration caused by cardiotoxin injury (**Fig. 5A & Suppl. Fig. 11**). Immunofluorescence of MP marker MyoD and proliferation marker bromodeoxyuridine (Brdu), revealed significantly more proliferating MPs in p107KO tibialis anterior (TA) muscle compared to wild type regenerating muscle, 2 days post cardiotoxin injury (**Fig. 5A**).

**Fig 5.**
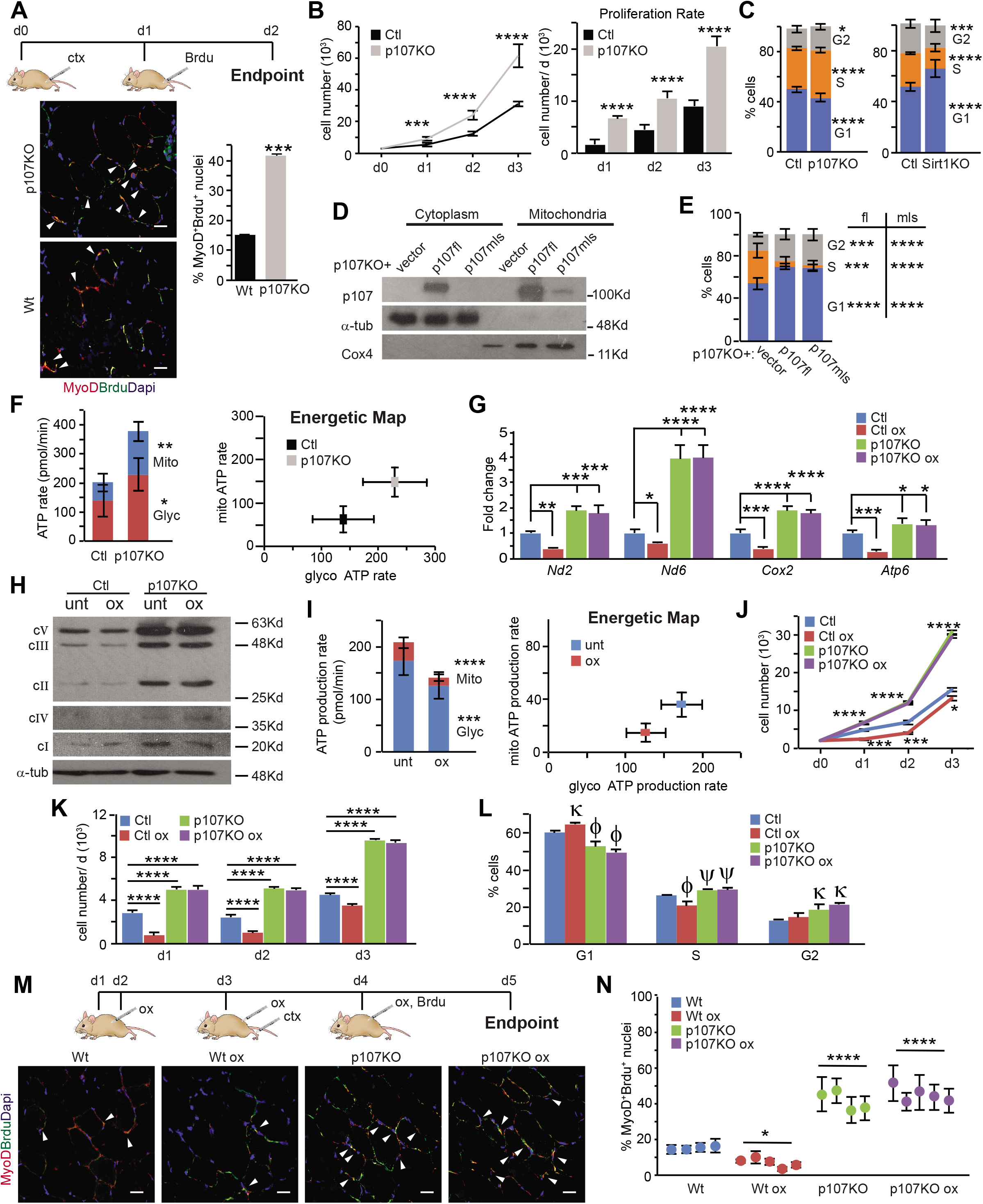
p107 directs the cell cycle rate through management of Oxphos generation. **(A)** Time course of treatments for wild type (Wt) and p107KO mice injected with cardiotoxin (ctx) and bromodeoxyuridine (Brdu). Representative confocal immunofluorescence merge image of MyoD, Brdu and Dapi and graph depicting the percentage of proliferating myogenic progenitors from tibialis anterior muscle at day 2 post injury, n=4, \*\*\**p*<0.001; Student T-test. Arrows denote Brdu and MyoD positive nuclei. (scale bar 20μm) **(B)** Growth curve and proliferation rate during 3 days for control (Ctl) and p107KO c2MPs, n=6-8; \*\*\**p*<0.001, \*\*\*\**p*<0.0001; Student T-test. **(C)** Cell cycle analysis by flow cytometry for Ctl, p107KO and Sirt1KO c2MPs, n=6-9; \**p*<0.05; \*\*\**p*<0.001 \*\*\*\**p*<0.0001; Student T-test. **(D)** Representative Western blot of cytoplasmic (Cyto) and mitochondria (Mito) fractions of p107KO c2MPs that were transfected with empty vector alone or with full length p107 (p107fl) or mitochondria localized p107 (p107mls). **(E)** Cell cycle analysis by flow cytometry for cells in (D), n=6-9; \*\*\**p*<0.001, \*\*\*\**p*<0.0001; Student T-test. **(F)** Live cell metabolic analysis of ATP production rate from mitochondria (Mito) and glycolysis (Glyc) and energetic map for Ctl and p107KO c2MPs, n=4-5; \**p*<0.05, \*\**p*<0.01; two-way Anova and post hoc Tukey. **(G)** qPCR of *Nd2, Nd6, Cox2* and *Atp6* for Ctl and p107KO c2MPs in the presence or absence of oxamate (ox), n=4. \**p*<0.01, \*\**p*<0.01, \*\*\**p*<0.001, \*\*\*\**p*<0.0001; two-way Anova and post hoc Tukey. **(H)** Representative Western blot for Ctl and p107KO c2MPs untreated or treated with ox for electron transport chain complex. **(I)** Live cell metabolic analysis for ATP production of c2MPs untreated or treated with ox, n=6-8. **(J)** Growth curve, n=6-8, **(K)** proliferation rate, n=6-8, **(L)** Cell cycle analysis, n=6-8 for Ctl and p107KO c2MPs untreated or treated with ox; \*\*\**p*<0.001, \*\*\*\**p*<0.0001 and ψ<0.05, k<0.001, ϕ<0.0001. **(M)** Time course treatments for Wt and p107KO mice injected with ox, ctx and Brdu and representative confocal immunofluorescence merge of MyoD, Brduand Dapi (scale bar 20μm). Arrows denote Brdu and MyoD positive nuclei and **(N**) graph depicting the percentage of proliferating MPs from tibialis anterior muscle at day 2 post injury, n=5; \**p*<0.05, \*\*\*\**p*<0.001; two-way Anova with post hoc Tukey.

We confirmed that the proliferative differences in the p107 genetically deleted mice compared to wild type littermates were cell autonomous by considering control and p107KO c2MPs. By counting cells over several days and performing flow cytometry cell cycle analysis, we found p107KO c2MPs had almost twice the cell cycle rate, with significantly increased cell numbers in S-phase compared to control c2MPs, as had been previously shown with p107KO prMPs (LeCouter *et al*, 1998) (**Fig. 5B & 5C**). Contrarily, Sirt1KO cells that had increased levels of p107 in the mitochondria (**Fig. 4B**) exhibited significantly decelerated cell cycle progression with significantly fewer cells in S-phase compared to controls **(Fig. 5C)**. We assessed if the mitochondrial function of p107 was essential to the cell cycle rate reduction, by restoring p107 mitochondrial localization in p107KO cells. This was accomplished by over expressing p107fl or p107 containing a mitochondrial localization sequence (p107mls) (**Fig. 5D & Suppl. Fig. 12A**). Restoring p107 to the genetically deleted cells significantly inhibited cell cycle progression by blocking cells in G1 phase of the cell cycle, which corresponded to reduced mitochondrial gene expression (**Fig. 5E & Suppl. Fig. 12B**). This suggests that p107 might direct the cell cycle rate through management of Oxphos generation.

We tested this possibility by using live cell metabolic imaging with Seahorse, which showed that the augmented cell cycle rate in p107KO cells compared to controls corresponded to a significant increase rate of ATP generation from Oxphos and glycolysis, including an increase in the mitochondrial to glycolytic ATP production rate ratio (**Fig. 5F)**. In contrast, live cell metabolic imaging by Seahorse showed that p107KO c2MPs that over-expressed p107fl or p107mls, which is directed to the mitochondria, resulted in significant attenuation of ATP generation capacity **(Suppl. Fig. 13)**.

We next appraised if modulation of p107 mitochondrial function to control ATP production might influence MP proliferation. For this, the cytoplasmic NAD^+^/NADH ratio was manipulated with addition of ox (**Fig. 3I**). As ox inhibits Ldha, it results in glycolytic flux relying almost entirely on NAD^+^ generated by Oxphos. It also provides increased NADH, which augmented p107 mitochondrial localization and decreased mtDNA gene expression (**Fig. 3J, 3k & 5G**). However, ox did not affect mitochondrial gene expression in p107KO cells (**Fig. 5G**). Moreover, Western blotting revealed that ox treatment also decreased ETC complex levels in c2MPs but showed no change in p107KO c2MPs (**Fig. 5H**).

Live metabolic imaging of c2MPs demonstrated that the reduction in ETC complexes with ox was associated with a significant decrease in ATP generation from Oxphos as well as glycolysis (**Fig. 5I and Suppl. Fig. 14**). The reduced ATP production was coupled to a significant decrease in cell growth and proliferation rate (**Fig. 5J & 5K**) as well as the cell cycle rate and S-phase (**Fig 5L**). However, ox treatment of p107KO c2MPs did not alter ATP levels, and the cell cycle rate (**Fig. 5I, 5J, 5K & 5L**). Taken together, these results suggest that Oxphos regulation by p107 controls c2MP proliferative rates.

We confirmed these findings in vivo by treatment of mice with ox during the regeneration process. Wild type mice treated with ox showed significantly fewer proliferating MyoD^+^Brdu^+^ MPs compared to control untreated mice (**Fig. 5M, 5N & Suppl. Fig. 15**). In contrast, the muscles from p107KO mice treated or untreated with ox showed no difference in proliferating MPs (**Fig. 5M, 5N and Suppl. Fig. 15**).

Collectively, these results suggest that p107 mitochondrial function controls cell cycle rate by regulating ETC complex formation in a Sirt1 dependent manner (Suppl. Fig. 16).

## Discussion

Our results for the first time support an unanticipated mitochondrial function for p107 as a regulator of ATP generation. We used a series of complimentary approaches that underscore p107 operation within the mitochondria as a transcriptional co-repressor. Biochemical mitochondrial fractionation showed that it is located in the matrix and interacts at the D-loop promoter of mtDNA to repress gene expression. By controlling gene expression in this manner, we found that p107 directly impacted the Oxphos capacity of MP cells and restricted their proliferation rate. Accordingly, these findings showcase that p107 arbitrates cell cycle rate through metabolic control. Indeed, p107 mitochondrial function is controlled by the cellular redox status and in particular the energy sensor Sirt1 an NAD^+^ dependent deacetylase. Intriguingly, as p107 is almost always shown to be expressed only in proliferating cells, the p107-Sirt1 pathway in MPs is possibly a universal cell mechanism utilized during division.

For many years, assigning a functional role for p107 has been ambiguous, unlike family member Rb that is a bona fide G1 phase restriction check point factor(Wirt & Sage, 2010). In this study we found that p107 acts as a metabolic checkpoint molecule for cellular energy status, placing it within the pantheon of other well-known nutrient sensing cell growth checkpoint molecules(Foster *et al*, 2010). During cell division, it is unclear how glycolysis and Oxphos collectively operate to regulate the yield of ATP. Our data show a major rationale for the ubiquitous expression of p107 in proliferating cells as part of a mechanism involved in this interplay. Our results highlight that p107 influences ATP production in the mitochondria based on the energy generated in the cytoplasm, in a Sirt1 dependent manner. The lowering of the NAD^+^/NADH ratio by a higher glycolytic flux was concomitant with a decreased capacity to generate ATP from Oxphos. We believe this is in part due to increasing levels of p107 interacting at the mtDNA where it represses gene expression to limit ETC complex formation. At this time, it is not known how p107 is shuttled across the outer and inner mitochondrial membranes. It might be a result of post-translational modification by acetylation and/or phosphorylation.

The role of p107 in repressing mtDNA transcription might ensure that ATP levels remain compatible with the demands of the proliferating cells and might also help steer the TCA cycle away from producing reducing equivalents in favour of biomass production(Vander Heiden & DeBerardinis, 2017). Notably, p107 is found to relocate from the cytoplasm to nucleus at late G1 or G1/S phase(Lindeman *et al*, 1997; Muller *et al*, 1997; Rodier *et al*, 2005; Verona *et al*, 1997; Zini *et al*, 2001). This is when a glycolytic to Oxphos switch occurs and when it silences Ldha and Pdk2 gene expression on nuclear promoters to further increase Oxphos capacity as shown in adipocyte progenitors(Porras *et al*., 2017). Thus, we propose that p107 dynamically modulates cell metabolism during the cell cycle through promoter repression in two different organelles. Through inhibiting Oxphos in G1, dependent on the nutrient load, and increasing Oxphos in S phase p107 is able to exert dual influences on cell cycle progression.

The high rate of ATP generated by Oxphos and glycolysis in p107KO MPs can explain the accelerated cell cycle rate. The higher proliferative rate is also characteristic of p107KO MEFs and primary MPs(LeCouter *et al*., 1998) and was evidenced in the significantly greater number of proliferating MPs within regenerating p107KO skeletal muscle. Conversely, if p107 function is dysregulated and forced to remain in the mitochondria, the reduced ATP generation capacity alarmingly decreases cell cycle efficiency. Thus, the decreased proliferative capacity of p107 over expressing cells found in many reports might now be attributed to a mitochondrial role in repressing mitochondrial gene expression, which is required for G1 cell cycle progression. Indeed, over expression of p107 can inhibit cell proliferation and arrest cell cycle at G1 in many cell types(Leng *et al*, 2002; Wirt & Sage, 2010; Zhu *et al*, 1993) and delay the G1 to S phase entry in rat fibroblast cell lines(Rodier *et al*., 2005).

Our data strongly support the concept that the NAD^+^/NADH ratio controls p107 function through Sirt1, which is activated by NAD^+^. Moreover, we and others have shown that Sirt1 regulation of SC and MP cell cycle parallels p107 mitochondrial control of proliferation. Indeed, Sirt1 has been shown to promote MP proliferation(Rathbone *et al*, 2009; Wang *et al*, 2016), which corresponds to p107 relocation from the mitochondria. Consistent with this idea, mice with activated Sirt1 (by caloric restriction or by NAD^+^ repletion) exhibited greater SC self-renewal and more proliferative MPs(Cerletti *et al*, 2012; Fulco *et al*, 2008; Ryall *et al*., 2015; Zhang *et al*., 2016). Conversely, primary activated SCs isolated from Sirt1KO, as well as conditional Sirt1KO, that would increase p107 mitochondrial localization and repress mtDNA gene expression, showed significantly lowered cell cycle as measured by the incorporation of nucleotide base analogs into DNA(Myers *et al*, 2019; Tang & Rando, 2014). Though the use of Sirt1KO cells confirmed that p107 functions through a Sirt1 dependent mechanism, it is unlikely that the functional interaction between p107 and Sirt1 is maintained during myogenic differentiation, as Sirt1 has a non-metabolic role during differentiation(Fulco *et al*, 2003; Ryall *et al*., 2015). Nonetheless, these findings set the stage for future evaluation of the mitochondrial function of p107 during SC activation, self-renewal and commitment.

In summary, our findings establish that a cell cycle regulator, p107, functions as a key and fundamental component of the cellular metabolism network during cell division. Indeed, it directly manipulates the energy generation capacity of mitochondria by indirectly sensing the glycolytic energy production through Sirt1. These results provide a conceptual advance for how proliferating cells regulate energy generation through the interplay between glycolysis and Oxphos. Importantly because of the ubiquitous p107 protein expression in most dividing cells, the findings identify a potential universal cellular mechanism with immense implications for studies on cancer cell proliferation and stem cell fate decisions.

## Materials and Methods

### Cells and mice

The c2c12 myogenic progenitor cell (c2MP) line was purchased from the American Tissue Type Culture (ATTC) and grown in Dulbecco’s Modified Eagle Medium (DMEM) containing 25mM glucose supplemented with 10% fetal bovine serum (FBS) and 1% penicillin streptomycin. All animal experiments were performed following the guidelines approved by the Animal Care Committee of York University. For derivation of prMPs and tissue immunofluorescence wild type and p107KO mice were from M. Rudnicki (LeCouter *et al*., 1998) maintained on a mixed (NMRI, C57/Bl6, FVB/N) background (Chen *et al*, 2004). The prMPs were isolated from single fibers obtained from the extensor digitoris longus muscles that were grown for 5 days before the media containing the prMPs was transferred to collagen coated fresh tissue culture plates. The prMPs were grown in Ham’s F10 Nutrient Mix Media supplemented with 20% FBS, 1% penicillin streptomycin and 2.5ng/ml bFGF (Peprotech).

For the nutrient specific experiments, c2MPs or prMPs were treated with glucose, galactose or glutamine. For drug specific treatment, cells were treated with oxamate, dichloroacetic acid, nicotinamide or resveratrol (for detailed protocol see Supplemental Materials and Methods). Western blot, nuclear and cytoplasmic extraction, growth curve and proliferation rate, see Supplemental Materials and Methods.

### Cloning

The p107mls expression plasmid that expresses p107 only in the mitochondria was made by cloning full length p107 into the pCMV6-OCT-HA-eGFP expression plasmid vector (Newman *et al*, 2016) that contains a mitochondrial localization signal. We used the following forward 5’-CACCAATTGATGTTCGAGGACAAGCCCCAC-3’ and reverse 5’-CACAAGCTTTTAATGATTTGCTCTTTCACT-3’ primer sets that contain the restriction sites Mfe1 and HindIII, respectively, to amplify full a length p107 insert from a p107 Ha tagged plasmid (Iwahori *et al*, 2017). The restriction enzyme digested full length p107 insert was then ligated to an EcoRI/HindIII digest of pCMV6-OCT-HA-eGFP, which removed the HA-eGFP sequences but retained the n-terminal mitochondrial localization signal (OCT). The calcium chloride method was used for transfections (see Supplemental Materials and Methods).

For overexpression studies, at least 4 different p107KO c2MPs were transfected as above with GFP mitochondrial localization empty vector pCMV6-OCT-HA-eGFP, (Newman *et al*., 2016), p107fl expressing full length p107 tagged HA, and p107mls expressing full length p107, which is directed to the mitochondria. For Sirt1 overexpression experiments, c2MPs were transfected with p107fl alone or with full length (Sirt1fl) or dominant negative (Sirt1dn) Sirt1 (Boily *et al*, 2009).

### p107KO and SirtKO cell lines derivation

Crspr/Cas9 was used to generate p107 and Sirt1 genetically deleted c2MP (p107KO and Sirt1KO) cell lines. For p107KO c2MPs, c2c12 cells were simultaneously transfected with 3 pLenti-U6-sgRNA-SFFV-Cas9-2A-Puro plasmids each containing a different sgRNA to target p107 sequences 110 CGTGAAGTCATCCAGGGCTT, 156 GGGAGAAGTTATACACTGGC and 350 AGTTTCGTGAGCGGATAGAA (Applied Biological Materials), and for Sirt1KO with 2 Double Nickase plasmids each containing a different sgRNA to target sequences 148 CGGACGAGCCGCTCCGCAAG and 110 CCATGGCGGCCGCCGCGGAA (Santa Cruz Biotechnology). For control cells, c2MPs cells were transfected by empty pLenti-U6-sgRNA-SFFV-Cas9-2A-Puro (Applied Biological Materials). See Supplemental Materials and Methods for detailed protocol.

### Cardiotoxin-induced muscle regeneration

Three-month-old anesthetized wild type and p107KO mice were injected intramuscularly in the tibialis anterior (TA) muscle with 40μl of cardiotoxin (ctx) Latoxan (Sigma) that was prepared by dissolving in water to a final concentration of 10μM. A day after ctx injury, bromodeoxyuridine (Brdu) at 100mg/kg was injected intraperitonially and TA muscles were collected on day 2 post ctx injection. Mice were also untreated or treated with 750mg/kg ox for four consecutive days, with ctx on the third day and Brdu on the fourth day, before the TA muscles were dissected on the fifth day for freezing.

### Mitochondrial isolation

Cells were washed in PBS, pelleted, dissolved in 5 times the packed volume with isolation buffer (0.25M Sucrose, 0.1% BSA, 0.2mM EDTA, 10mM HEPES with 1mg/ml of each pepstatin, leupeptin and aprotinin protease inhibitors), and homogenized in Dounce homogenizor on ice. The homogenate was centrifuged at 1000g at 4°C for 10 minutes. The supernatant was then centrifuged at 14000g for 15 min at 4°C and the resulting supernatant was saved as “cytosolic fraction”. The pellet representing the “mitochondrial fraction” was washed twice, dissolved in isolation buffer. The mitochondria were lysed by repeated freeze-thaw cycleson dry ice. Mitochondrial fractions were isolated using a hypotonic osmotic shock approach (Lu *et al*., 2009) (detailed protocol in Supplemental Materials and Methods.

### Mitochondrial and nuclear DNA content

To obtain the mitochondrial to nuclear DNA ratio (mtDNA/nDNA), cells grown on a 6cm tissue culture plate were untreated or treated with 1mM 5-aminoimidazole-4-carboxamide-1-β-D-ribofuranoside (AICAR) (Toronto Research Chemicals) in presence of 5.5mM or 25mM glucose, with or without 10mM Nam for 24 hours. (detailed protocol in Supplemental Materials and Methods.

### Primary antibodies used

α-tubulin (66031, Proteintech); Cox4 (ab16056, Abcam) and Total OXPHOS rodent WB antibody cocktail (Abcam); p107-C18, p107-SD9, histone H3-C16, Sirt1-B7, IgG-D7, Ha-tag-F7, Brdu-MoBU-1, Pax7-EE8 (Santa Cruz Biotech); MyoD (Novus Biologicals) and Sirt1-D1D7 (Cell Signaling).

### Co-immunoprecipitaion assay

For Immunoprecipitation (IP), protein lysates were pre-cleared with 50μl protein A/G plus agarose beads (Santa Cruz Biotechnology) by rocking at 4°C for an hour. The sample was centrifuged at 15000 rpm for a minute. Fresh protein A/G agarose beads along with 5μg of p107-C18, Sirt1-B7 or IgG-D7 antibody were added to the supernatant and rocked overnight at 4°C. The next day the pellets were washed 3 times with wash buffer (50mM HEPES pH 7.0, 250mM NaCl and 0.1% Np-40) and loaded onto polyacrylamide gels and Western blotted for p107-SD9 or Sirt1-D1D7. Inputs represent 10% of lysates that were immunoprecipitated.

### qPCR analysis

qPCR experiments were performed according to the MIQE (Minimum Information for Publication of Quantitative Real-Time PCR Experiments) guidelines (Bustin et al., 2009). (Supplemental Materials and Methods) and for primer sets used (Suppl. Table 2).

### Quantitative chromatin immunoprecipitation assay (qChIP)

qChIP was performed according to De Souza et al (De Sousa *et al*., 2014), (detailed protocol in Supplemental Materials and Methods). Relative occupancy was determined by amplifying isolated DNA fragments using the D-loop primers (**Suppl. Table 2**) and analysed using the ΔΔCt method.

### NAD^+^/NADH and ATP generation assays

NAD^+^/NADH assay was performed as previously published (Porras *et al*., 2017), view detailed protocol in Supplemental materials and methods. ATP production capacity of isolated mitochondria was measured using an ATP determination kit (ThermoFisher Scientific, USA) as per manufacturer’s instructions (Supplemental Materials and Methods). ATP production for each sample was normalized to total protein using Bradford assay kit (BioBasic).

### Immunocytochemistry and confocal imaging

Cells were washed in PBS, fixed for 5 minutes with 95% methanol and permeabilized for 30 minutes at 4°C with blocking buffer (3% BSA and 0.1% saponin in PBS). Then incubated with primary antibodies followed by secondary antibodies 1 hour each, with intermittent washing Finally, cells were incubated in 4’,6-diamidino-2-phenylindole (Dapi) and confocal images and Z-stacks were obtained using the Axio Observer.Z1 microscope with alpha Plan-Apochromat 63x/Oil DIC (UV) M27 (Zeiss) using Axiocam MR R3. (See supplemental Materials and Methods for detail).

### Mitochondrial length and area measurement

Live cell imaging by Axio Observer.Z1 (Zeiss) microscope with alpha plan-apochromat 40x/Oil DIC (UV) M27 (Zeiss) in an environment chamber (5% CO_2_; 37°C) using Axiocam MR R3 (Zeiss). **C**ells were stained with MitoView red (Biotium) and the mitochondrial length and area were measured by using Image J software (detailed protocol in Supplemental Materials and Methods).

### Immunohistochemistry

Frozen muscle tissue samples were fixed with 4% paraformaldehyde for 15 minutes and for antigen retrieval 2N HCl was added for 20 minutes followed by 40mM sodium citrate. After washing in PBS, muscle sections were blocked in blocking buffer (5% goat serum, 0.1% Triton X in PBS) for 30 minutes. The sections were then incubated with primary antibodies followed by secondary antibodies with intermittent washes. Finally, Dapi was added and the sections were imaged using confocal microscopy with the Axio Observer.Z1 microscope with alpha Plan-Apochromat 40x/Oil DIC (UV) M27 (Zeiss). See Supplemental Materials and Methods for detail.

### Flow cytometry

For cell cycle analysis 50000, Ctl and p107KO cells were treated or untreated with 2.5mM ox for 40 hours or Ctl and Sirt1 KO cells were grown in 5.5mM glucose, to activate Sirt1, for 20 hours or p107KO cells 24 hours post transfection with pCMV6-OCT-HA-eGFP alone or together with p107fl or p107mls, were used. Following fixation and washing, the cells were incubated in 50μg/ml propidium iodide (ThermoFisher Scientific) and 25μg/ml RNAse (Thermo Fisher Scientific) before loading on the Attune Nxt Flow Cytometer (Thermo Fisher Scientific). See Supplementary Materials and Methods for detailed protocol.

### Live cell ATP analysis (Seahorse)

3000 control or p107KO cells or p107KO cells 24 hours post transfection with pCMV6-OCT-HA-eGFP alone or together with p107fl or p107mls were seeded in DMEM (Wisent) containing 10% FBS and 1% penicillin streptomycin on microplates (Agilent Technologies) and treated according to the required experiment. For analysis, cells were washed in XF assay media supplemented with 10mM glucose, 1mM pyruvate and 2mM glutamine (Agilent Technologies) and assessed using the Seahorse XF real-time ATP rate assay kit (Agilent Technologies) on a Seahorse XFe96 extracellular flux analyzer (Agilent Technologies), with addition of 1.5μM of oligomycin and 0.5μM rotenone + antimycin A as per the manufacturer’s direction. The energy flux data in real time was determined using Wave 2.6 software (Agilent Technologies).

### Statistical analysis

Statistical analysis was performed by GraphPad Prism. Student’s t-tests were used unless otherwise stated. Results were considered to be statistically significant when *p* <0.05. Specific data was analyzed using an appropriate one-way or two-way analysis of variance (ANOVA) with a criterion of *p*<0.05. All significant differences for ANOVA testing were evaluated using a Tukey post hoc test. All data are mean ± SD.

## Acknowledgements

The authors thank Drs. Mireille Khacho and Tara Haas for critical insights and Dr. Haas for careful reading of the manuscript. Drs. Magdalena Jaklewicz and Geetika Phukan and Mayoorey Murigathasan for technical support. A.S. is a recipient of a Natural Sciences and Engineering Research Council of Canada research grant (RGPIN-2018-05937).

## Author Contributions

A.S. planned and managed the research activity, wrote the original draft and acquired financial support for the project D.B. and A.S. critically appraised and reviewed the draft, made the figures, formulated the ideas, developed and designed the methodology, validated the research outputs, D.B., A.S. O.J.O conducted the experiments and formally analyzed the data

## References

Bhattacharya D, Scime A (2020) Mitochondrial Function in Muscle Stem Cell Fates. Front Cell Dev Biol 8: 480

Boily G, He XH, Pearce B, Jardine K, McBurney MW (2009) SirT1-null mice develop tumors at normal rates but are poorly protected by resveratrol. Oncogene 28: 2882–2893

Canto C, Menzies KJ, Auwerx J (2015) NAD(+) Metabolism and the Control of Energy Homeostasis: A Balancing Act between Mitochondria and the Nucleus. Cell Metab 22: 31–53

Cerletti M, Jang YC, Finley LW, Haigis MC, Wagers AJ (2012) Short-term calorie restriction enhances skeletal muscle stem cell function. Cell Stem Cell 10: 515–519

Cerutti R, Pirinen E, Lamperti C, Marchet S, Sauve AA, Li W, Leoni V, Schon EA, Dantzer F, Auwerx J et al (2014) NAD(+)-dependent activation of Sirt1 corrects the phenotype in a mouse model of mitochondrial disease. Cell Metab 19: 1042–1049

Chen D, Livne-bar I, Vanderluit JL, Slack RS, Agochiya M, Bremner R (2004) Cell-specific effects of RB or RB/p107 loss on retinal development implicate an intrinsically death-resistant cell-of-origin in retinoblastoma. Cancer Cell 5: 539–551

Dali-Youcef N, Mataki C, Coste A, Messaddeq N, Giroud S, Blanc S, Koehl C, Champy MF, Chambon P, Fajas L et al (2007) Adipose tissue-specific inactivation of the retinoblastoma protein protects against diabesity because of increased energy expenditure. Proc Natl Acad Sci U S A 104: 10703–10708

De Sousa M, Porras DP, Perry CG, Seale P, Scime A (2014) p107 is a crucial regulator for determining the adipocyte lineage fate choices of stem cells. Stem Cells 32: 1323–1336

Folmes CD, Nelson TJ, Dzeja PP, Terzic A (2012) Energy metabolism plasticity enables stemness programs. Ann N Y Acad Sci 1254: 82–89

Foster DA, Yellen P, Xu L, Saqcena M (2010) Regulation of G1 Cell Cycle Progression: Distinguishing the Restriction Point from a Nutrient-Sensing Cell Growth Checkpoint(s). Genes Cancer 1: 1124–1131

Fulco M, Cen Y, Zhao P, Hoffman EP, McBurney MW, Sauve AA, Sartorelli V (2008) Glucose restriction inhibits skeletal myoblast differentiation by activating SIRT1 through AMPK-mediated regulation of Nampt. Dev Cell 14: 661–673

Fulco M, Schiltz RL, Iezzi S, King MT, Zhao P, Kashiwaya Y, Hoffman E, Veech RL, Sartorelli V (2003) Sir2 regulates skeletal muscle differentiation as a potential sensor of the redox state. Molecular cell 12: 51–62

Ghosh-Choudhary S, Liu J, Finkel T (2020) Metabolic Regulation of Cell Fate and Function. Trends Cell Biol 30: 201–212

Gustafsson CM, Falkenberg M, Larsson NG (2016) Maintenance and Expression of Mammalian Mitochondrial DNA. Annu Rev Biochem 85: 133–160

Iwahori S, Umana AC, VanDeusen HR, Kalejta RF (2017) Human cytomegalovirus-encoded viral cyclin-dependent kinase (v-CDK) UL97 phosphorylates and inactivates the retinoblastoma protein-related p107 and p130 proteins. J Biol Chem 292: 6583–6599

Jones RA, Robinson TJ, Liu JC, Shrestha M, Voisin V, Ju Y, Chung PE, Pellecchia G, Fell VL, Bae S et al (2016) RB1 deficiency in triple-negative breast cancer induces mitochondrial protein translation. J Clin Invest 126: 3739–3757

Khacho M, Clark A, Svoboda DS, Azzi J, MacLaurin JG, Meghaizel C, Sesaki H, Lagace DC, Germain M, Harper ME et al (2016) Mitochondrial Dynamics Impacts Stem Cell Identity and Fate Decisions by Regulating a Nuclear Transcriptional Program. Cell Stem Cell 19: 232–247

Lapuente-Brun E, Moreno-Loshuertos R, Acin-Perez R, Latorre-Pellicer A, Colas C, Balsa E, Perales-Clemente E, Quiros PM, Calvo E, Rodriguez-Hernandez MA et al (2013) Supercomplex assembly determines electron flux in the mitochondrial electron transport chain. Science 340: 1567–1570

LeCouter JE, Kablar B, Hardy WR, Ying C, Megeney LA, May LL, Rudnicki MA (1998) Strain-dependent myeloid hyperplasia, growth deficiency, and accelerated cell cycle in mice lacking the Rb-related p107 gene. Mol Cell Biol 18: 7455–7465

Leng X, Noble M, Adams PD, Qin J, Harper JW (2002) Reversal of growth suppression by p107 via direct phosphorylation by cyclin D1/cyclin-dependent kinase 4. Mol Cell Biol 22: 2242–2254

Lindeman GJ, Gaubatz S, Livingston DM, Ginsberg D (1997) The subcellular localization of E2F-4 is cell-cycle dependent. Proc Natl Acad Sci U S A 94: 5095–5100

Locasale JW, Cantley LC (2011) Metabolic flux and the regulation of mammalian cell growth. Cell Metab 14: 443–451

Lu G, Sun H, Korge P, Koehler CM, Weiss JN, Wang Y (2009) Functional characterization of a mitochondrial Ser/Thr protein phosphatase in cell death regulation. Methods in enzymology 457: 255–273

Martinez-Reyes I, Chandel NS (2020) Mitochondrial TCA cycle metabolites control physiology and disease. Nature communications 11: 102

Martinez-Reyes I, Diebold LP, Kong H, Schieber M, Huang H, Hensley CT, Mehta MM, Wang T, Santos JH, Woychik R et al (2016) TCA Cycle and Mitochondrial Membrane Potential Are Necessary for Diverse Biological Functions. Molecular cell 61: 199–209

Mendelsohn BA, Bennett NK, Darch MA, Yu K, Nguyen MK, Pucciarelli D, Nelson M, Horlbeck MA, Gilbert LA, Hyun W et al (2018) A high-throughput screen of real-time ATP levels in individual cells reveals mechanisms of energy failure. PLoS Biol 16: e2004624

Mishra P, Carelli V, Manfredi G, Chan DC (2014) Proteolytic cleavage of Opa1 stimulates mitochondrial inner membrane fusion and couples fusion to oxidative phosphorylation. Cell Metab 19: 630–641

Mishra P, Chan DC (2016) Metabolic regulation of mitochondrial dynamics. J Cell Biol 212: 379–387

Mitra K, Wunder C, Roysam B, Lin G, Lippincott-Schwartz J (2009) A hyperfused mitochondrial state achieved at G1-S regulates cyclin E buildup and entry into S phase. Proc Natl Acad Sci U S A 106: 11960–11965

Muller H, Moroni MC, Vigo E, Petersen BO, Bartek J, Helin K (1997) Induction of S-phase entry by E2F transcription factors depends on their nuclear localization. Mol Cell Biol 17: 5508–5520

Myers MJ, Shepherd DL, Durr AJ, Stanton DS, Mohamed JS, Hollander JM, Alway SE (2019) The role of SIRT1 in skeletal muscle function and repair of older mice. J Cachexia Sarcopenia Muscle 10: 929–949

Newman LE, Schiavon C, Kahn RA (2016) Plasmids for variable expression of proteins targeted to the mitochondrial matrix or intermembrane space. Cell Logist 6: e1247939

Petrov PD, Ribot J, Lopez-Mejia IC, Fajas L, Palou A, Bonet ML (2016) Retinoblastoma Protein Knockdown Favors Oxidative Metabolism and Glucose and Fatty Acid Disposal in Muscle Cells. J Cell Physiol 231: 708–718

Porras DP, Abbaszadeh M, Bhattacharya D, D’Souza NC, Edjiu NR, Perry CGR, Scime A (2017) p107 Determines a Metabolic Checkpoint Required for Adipocyte Lineage Fates. Stem Cells 35: 1378–1391

Rathbone CR, Booth FW, Lees SJ (2009) Sirt1 increases skeletal muscle precursor cell proliferation. Eur J Cell Biol 88: 35–44

Rodier G, Makris C, Coulombe P, Scime A, Nakayama K, Nakayama KI, Meloche S (2005) p107 inhibits G1 to S phase progression by down-regulating expression of the F-box protein Skp2. J Cell Biol 168: 55–66

Ryall JG, Dell’Orso S, Derfoul A, Juan A, Zare H, Feng X, Clermont D, Koulnis M, Gutierrez-Cruz G, Fulco M et al (2015) The NAD(+)-dependent SIRT1 deacetylase translates a metabolic switch into regulatory epigenetics in skeletal muscle stem cells. Cell Stem Cell 16: 171–183

Salazar-Roa M, Malumbres M (2017) Fueling the Cell Division Cycle. Trends Cell Biol 27: 69–81

Scime A, Grenier G, Huh MS, Gillespie MA, Bevilacqua L, Harper ME, Rudnicki MA (2005) Rb and p107 regulate preadipocyte differentiation into white versus brown fat through repression of PGC-1alpha. Cell Metab 2: 283–295

Scime A, Soleimani VD, Bentzinger CF, Gillespie MA, Le Grand F, Grenier G, Bevilacqua L, Harper ME, Rudnicki MA (2010) Oxidative status of muscle is determined by p107 regulation of PGC-1a. J Cell Biol 190: 651–662

Tang AH, Rando TA (2014) Induction of autophagy supports the bioenergetic demands of quiescent muscle stem cell activation. The EMBO journal 33: 2782–2797

Vander Heiden MG, DeBerardinis RJ (2017) Understanding the Intersections between Metabolism and Cancer Biology. Cell 168: 657–669

Verona R, Moberg K, Estes S, Starz M, Vernon JP, Lees JA (1997) E2F activity is regulated by cell cycle-dependent changes in subcellular localization. Mol Cell Biol 17: 7268–7282

Wang L, Zhang T, Xi Y, Yang C, Sun C, Li D (2016) Sirtuin 1 promotes the proliferation of C2C12 myoblast cells via the myostatin signaling pathway. Mol Med Rep 14: 1309–1315

Wheaton WW, Weinberg SE, Hamanaka RB, Soberanes S, Sullivan LB, Anso E, Glasauer A, Dufour E, Mutlu GM, Budigner GS et al (2014) Metformin inhibits mitochondrial complex I of cancer cells to reduce tumorigenesis. Elife 3: e02242

Wirt SE, Sage J (2010) p107 in the public eye: an Rb understudy and more. Cell Div 5: 9

Zacksenhaus E, Shrestha M, Liu JC, Vorobieva I, Chung PED, Ju Y, Nir U, Jiang Z (2017) Mitochondrial OXPHOS Induced by RB1 Deficiency in Breast Cancer: Implications for Anabolic Metabolism, Stemness, and Metastasis. Trends Cancer 3: 768–779

Zhang H, Ryu D, Wu Y, Gariani K, Wang X, Luan P, D’Amico D, Ropelle ER, Lutolf MP, Aebersold R et al (2016) NAD(+) repletion improves mitochondrial and stem cell function and enhances life span in mice. Science 352: 1436–1443

Zhu L, van den Heuvel S, Helin K, Fattaey A, Ewen M, Livingston D, Dyson N, Harlow E (1993) Inhibition of cell proliferation by p107, a relative of the retinoblastoma protein. Genes Dev 7: 1111–1125

Zini N, Trimarchi C, Claudio PP, Stiegler P, Marinelli F, Maltarello MC, La Sala D, De Falco G, Russo G, Ammirati G et al (2001) pRb2/p130 and p107 control cell growth by multiple strategies and in association with different compartments within the nucleus. J Cell Physiol 189: 34–44

