## Supplemental Figures and Tables for "p107 mediated mitochondrial function controls muscle stem cell proliferative fates"

### Supplemental Figure 1

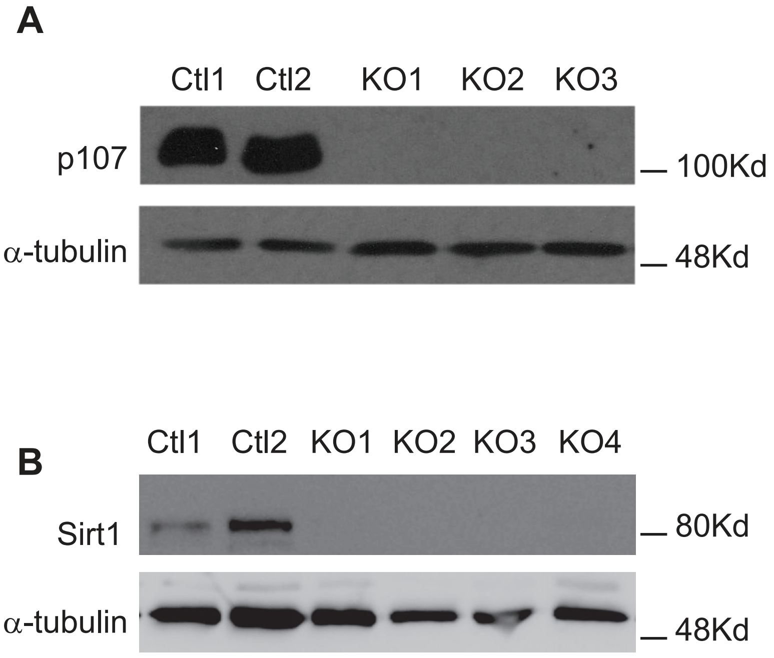


**Suppl. Fig. 1. Crispr/Cas9 generated p107 and Sirt1 genetically deleted c2MPs.** Representative Western blot for p107, Sirt1 and α-tubulin of different control c2MPs (Ctl) and **(A)** p107 or **(B)** Sirt1 genetically deleted c2MPs (KO).

### Supplemental Figure 2

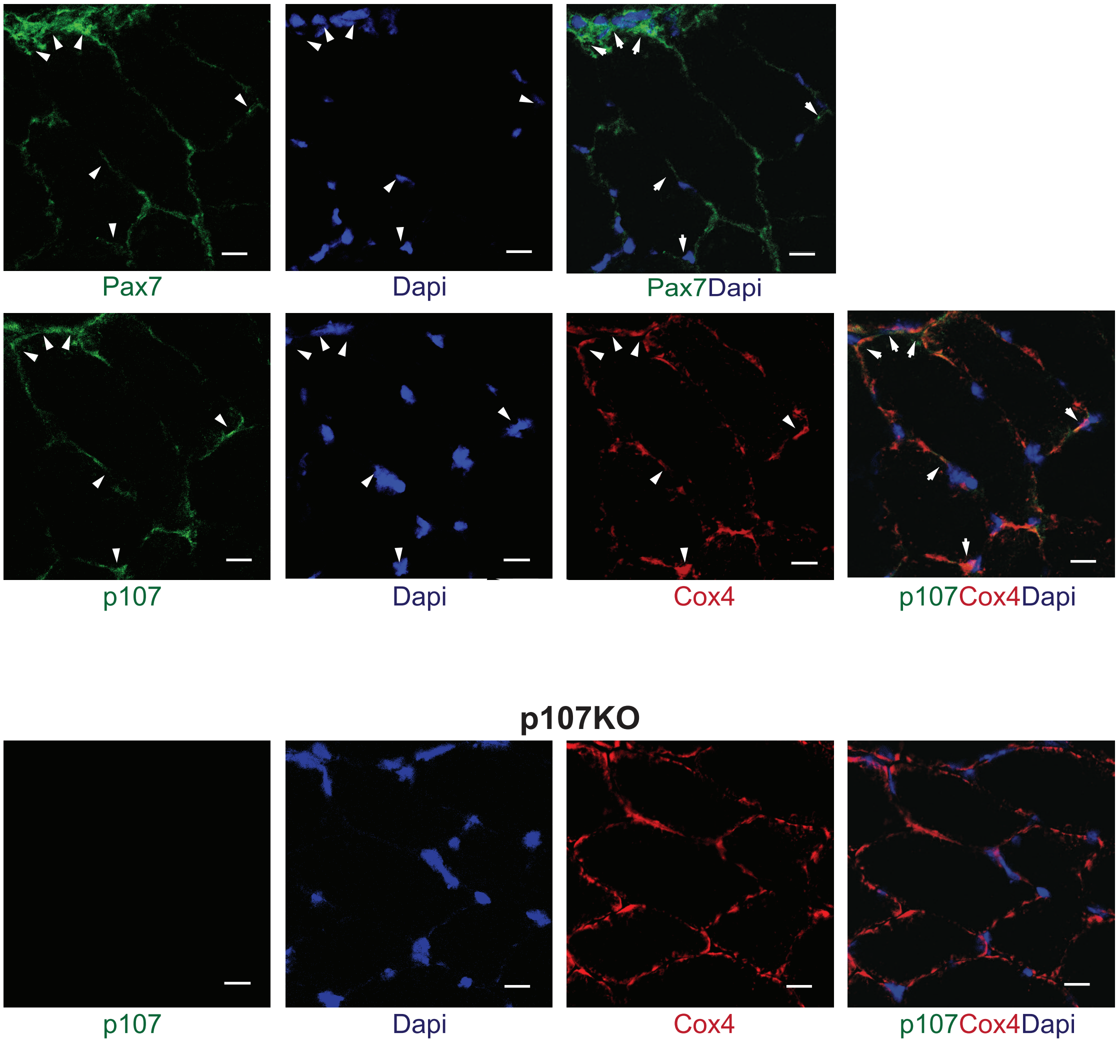


**Suppl. Fig. 2.** Confocal immunoflourescence microscopic image of wild type (Wt) tibialis anterior ( TA) muscle section 2 days post injury for Pax7 (green), Dapi (blue), Merge and p107 (green), Dapi (blue), Cox4 (red), Merge and of p107KO TA muscle section for p107 (green), Dapi (blue), Cox4 (red), Merge as a negative control (scale bar 20μm).

### Supplemental Figure 3

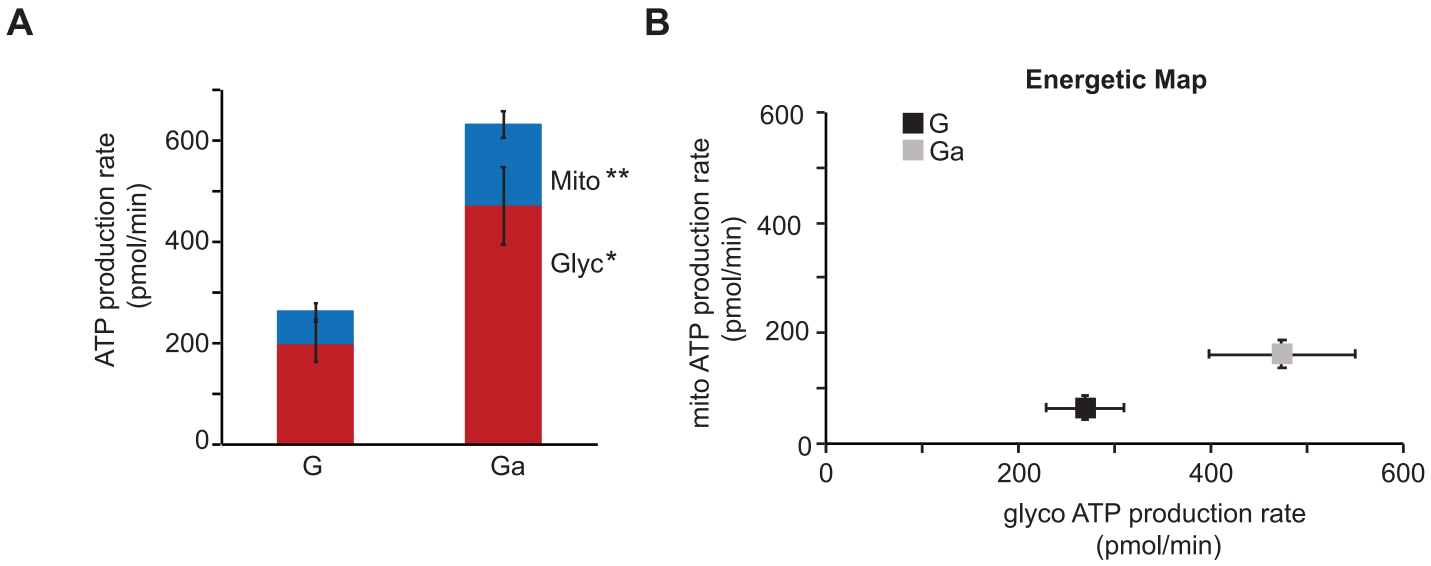


**Suppl. Fig. 3. Growth arrested cells have significantly enhanced production of total and mitochondrial ATP.** Live cell metabolic analysis by Seahorse of **(A)** ATP production rate from mitochondria (Mito) and glycolysis (Glyc) and **(B)** energetic map for proliferating (G) and growth arrested (Ga) c2MPs, n=6-8; asterisks denote significance, *$p<0.05$, **$p<0.01$; two-way Anova and post hoc Tukey.

#### Supplemental Figure 4

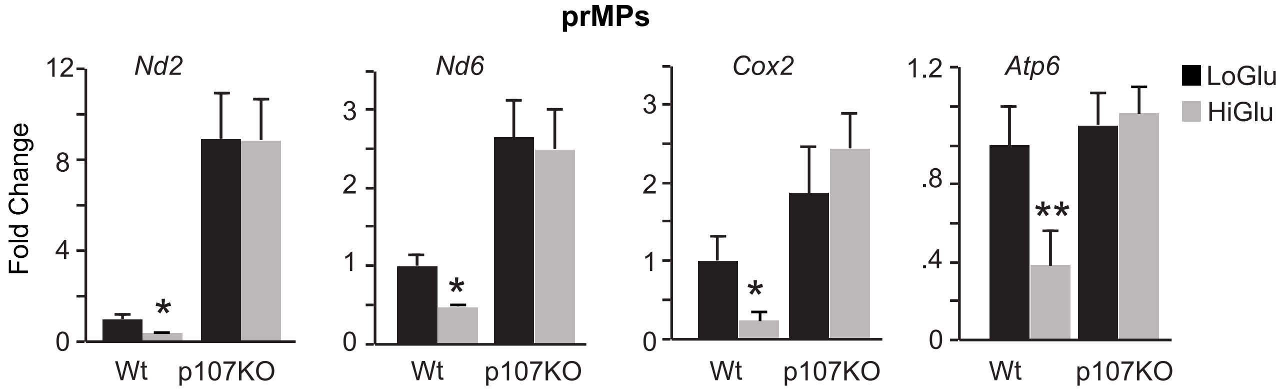


**Suppl. Fig. 4. Gene expression analysis of primary myogenic progenitors (prMPs) grown in media with low or high glucose concentrations.** Gene expression analysis by qPCR of mitochondrial encoded genes *Nd2*, *Nd6*, *Cox2* and *Atp6* from wild type (Wt) and p107 genetically deleted (p107KO) primary myogenic progenitors (prMPs) grown in stripped media containing 5.5mM (LoGlu) or 25mM (HiGlu) glucose, n=4, asterisks denote significance, *$p$<0.05 and **$p$<0.01; two-way Anova with post hoc Tukey.

### Supplemental Figure 5

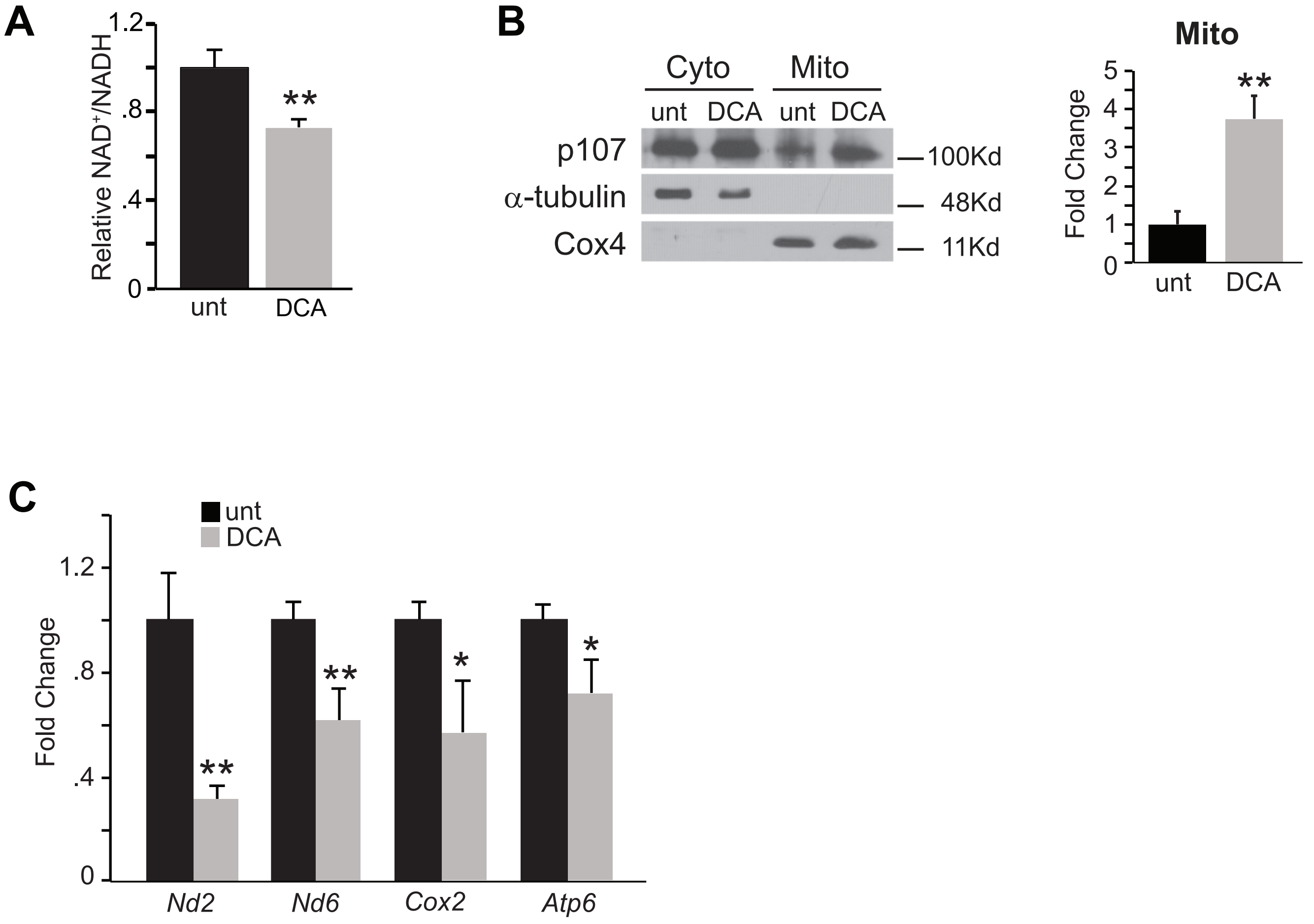


**Suppl. Fig. 5. NAD+/NADH regulation by dichloroacetic acid DCA influences p107 mitochondrial function. (A)** NAD+/NADH ratio for c2MPs untreated (unt) or treated with dichloroacetic acid (DCA). n=4, asterisks denote significance, $**p<0.01$; Student T test. **(B)** Representative Western blot and graphical representation of cytoplasmic (Cyto) and mitochondrial (Mito) fractions for p107, α-tubulin (cytoplasmic loading control) and Cox4 (mitochondria loading control) of cells in (A), n=3, asterisks denote significance, $**p<0.01$; Student T test. **(C)** qPCR analysis of cells in (A) for mitochondrial encoded genes *Nd2*, *Nd6*, *Cox2* and *Atp6*. n=3, asterisks denote significance, $*p<0.05$, $**p<0.01$; Student T test.

#### Supplemental Figure 6

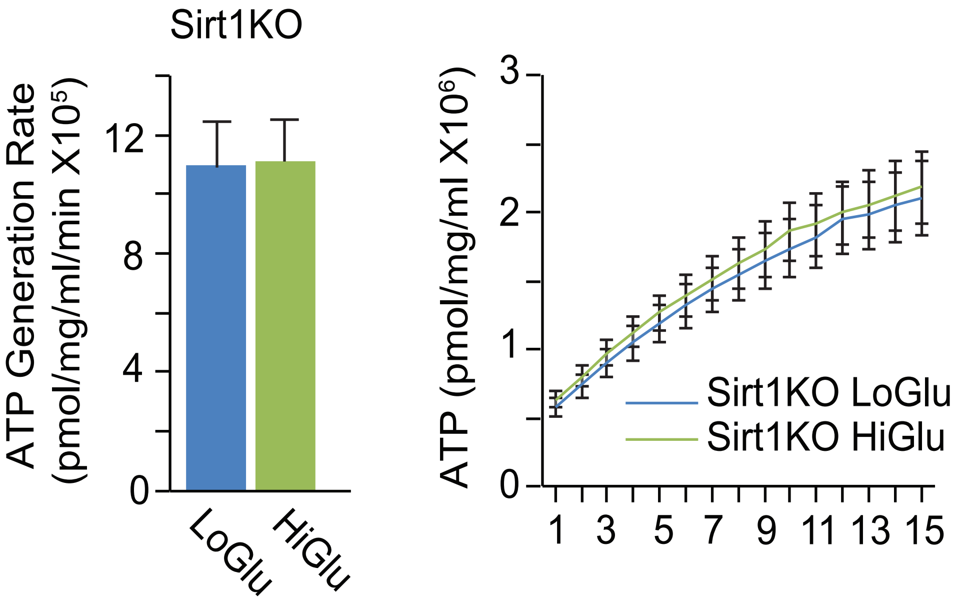


**Suppl. Fig. 6. Varying glucose concentrations do not influence ATP generation rate and capacity in isolated mitochondria of Sirt1KO cells.** Isolated mitochondrial ATP generation rate and capacity over time for Sirt1KO c2MPs grown in 5.5mM (LoGlu) and 25mM (HiGlu) glucose.

#### Supplemental Figure 7

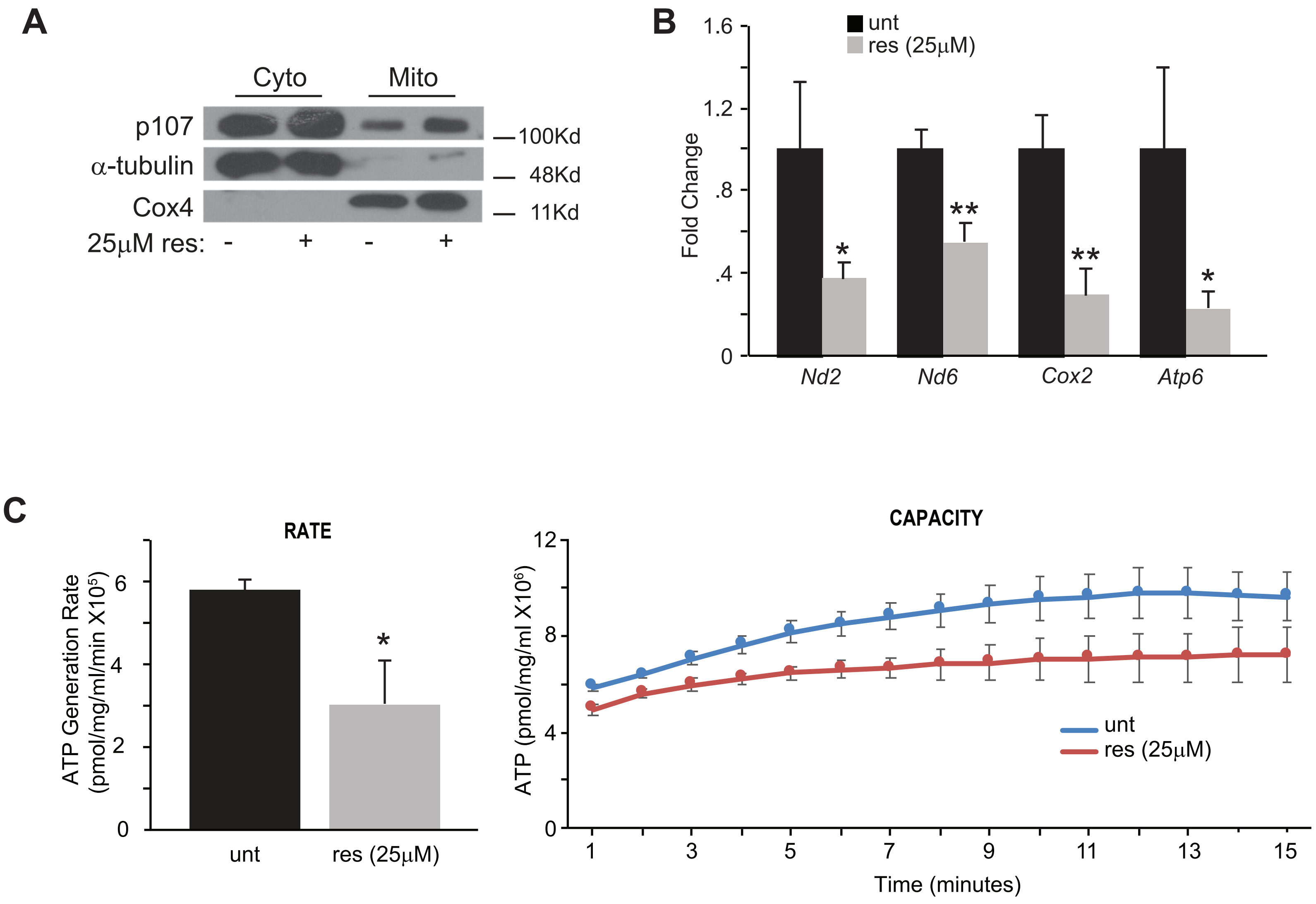


**Suppl. Fig. 7. A high resveratrol (25µM) concentration that inactivates Sirt1 increases p107 mitochondrial function. (A)** Representative Western blot of cytoplasmic (Cyto) and mitochondrial (Mito) fractions for p107, α-tubulin and Cox4 of c2MPs untreated (unt) or treated with a concentration (25µM) of resveratrol (res) that inactivates Sirt1. **(B)** Gene expression analysis by qPCR of mitochondrial encoded genes *Nd2*, *Nd6*, *Cox2* and *Atp6* for c2MPs untreated (unt) or treated with a concentration (25µM) of res that inactivates Sirt1, n=3-4; asterisks denote significance, *$p<0.05$, **$p<0.01$; Student T-test. **(C)** Isolated mitochondrial ATP generation rate and capacity over time for c2MPs untreated (unt) or treated with a concentration (25µM) of res that inactivates Sirt1; asterisks denote significance, *$p<0.05$, **$p<0.01$ ***$p<0.001$; Student T-test.

### Supplemental Figure 8

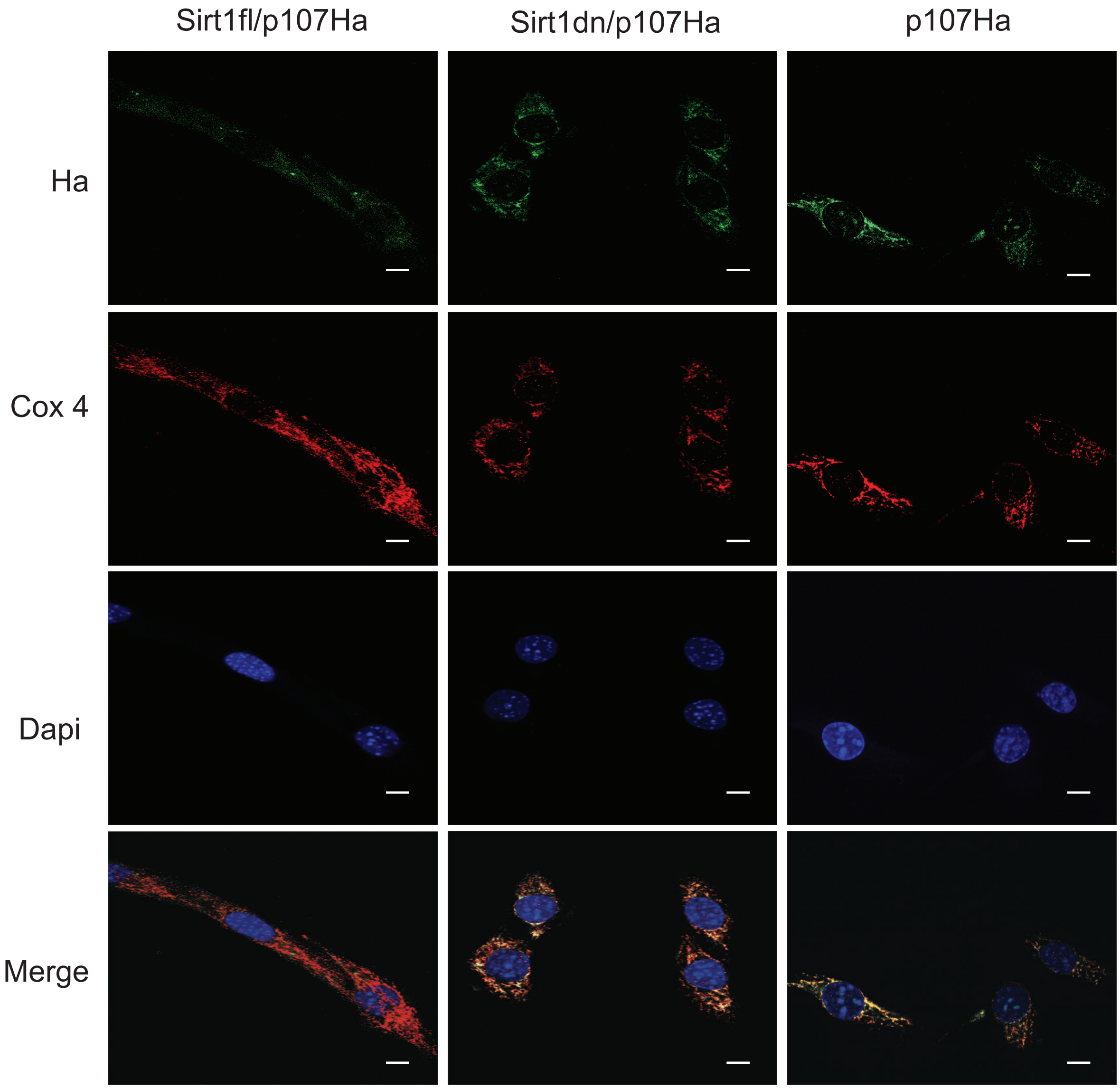


**Suppl. Fig. 8.** Confocal immunofluorescence microscopy for Ha (green), Cox4 (red), Dapi (blue) and Merge of c2MPs grown in stripped media containing 5.5mM glucose transfected with full length Ha-tagged p107 (p107Ha) alone or with full length Sirt1 (Sirt1fl) or dominant negative Sirt1 (Sirt1dn). (scale bar 10μm).

### Supplemental Figure 9

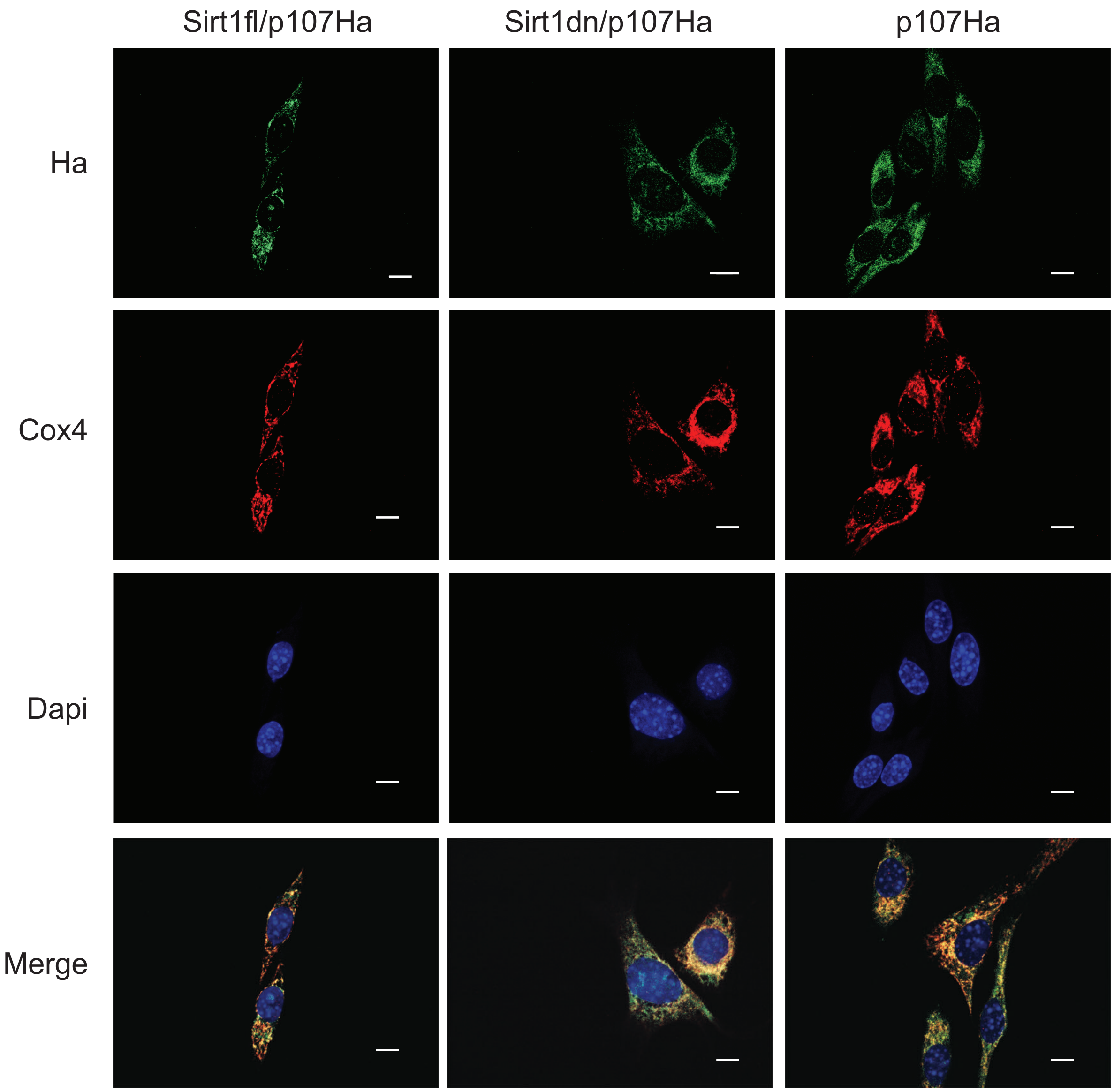


**Suppl. Fig. 9.** Confocal immunofluorescence microscopy for Ha (green), Cox4 (red), Dapi (blue) and Merge of c2MPs grown in stripped media containing 25mM glucose transfected with full length Ha-tagged p107 (p107Ha) alone or with full length Sirt1 (Sirt1fl) or dominant negative Sirt1 (Sirt1dn). (scale bar 10μm).

#### Supplemental Figure 10

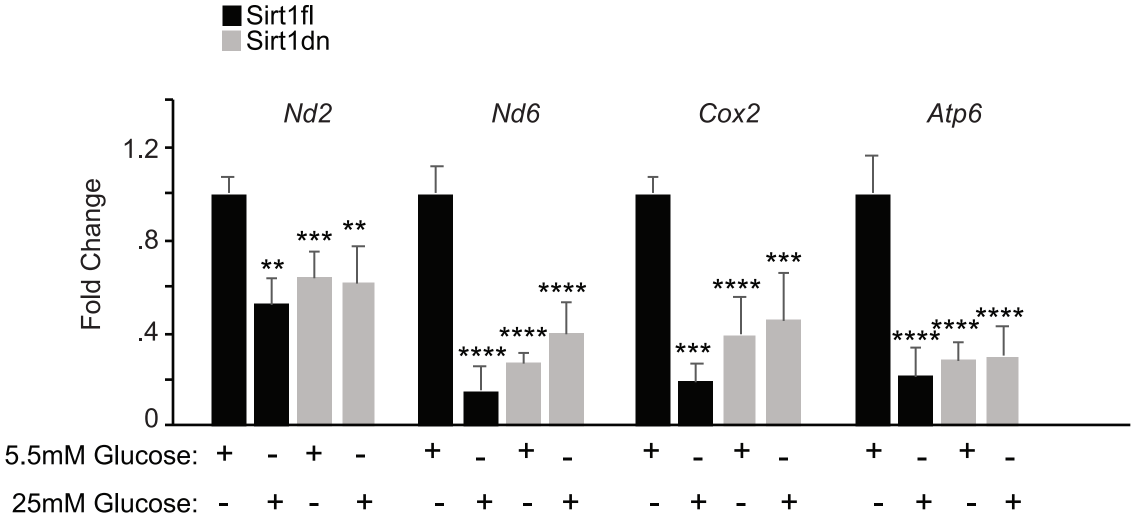


**Suppl. Fig. 10.** Gene expression analysis by qPCR of mitochondrial encoded genes *Nd2*, *Nd6*, *Cox2* and *Atp6* of c2MPs transfected with full length (Sirt1fl) or dominant negative (Sirt1dn) grown in stripped media containing 5.5mM or 25mM glucose n=4-5, asterisks denote significance, ***p*<0.01, ****p*<0.001, *****p*<0.0001; two-way Anova with post hoc Tukey.

### Supplemental Figure 11

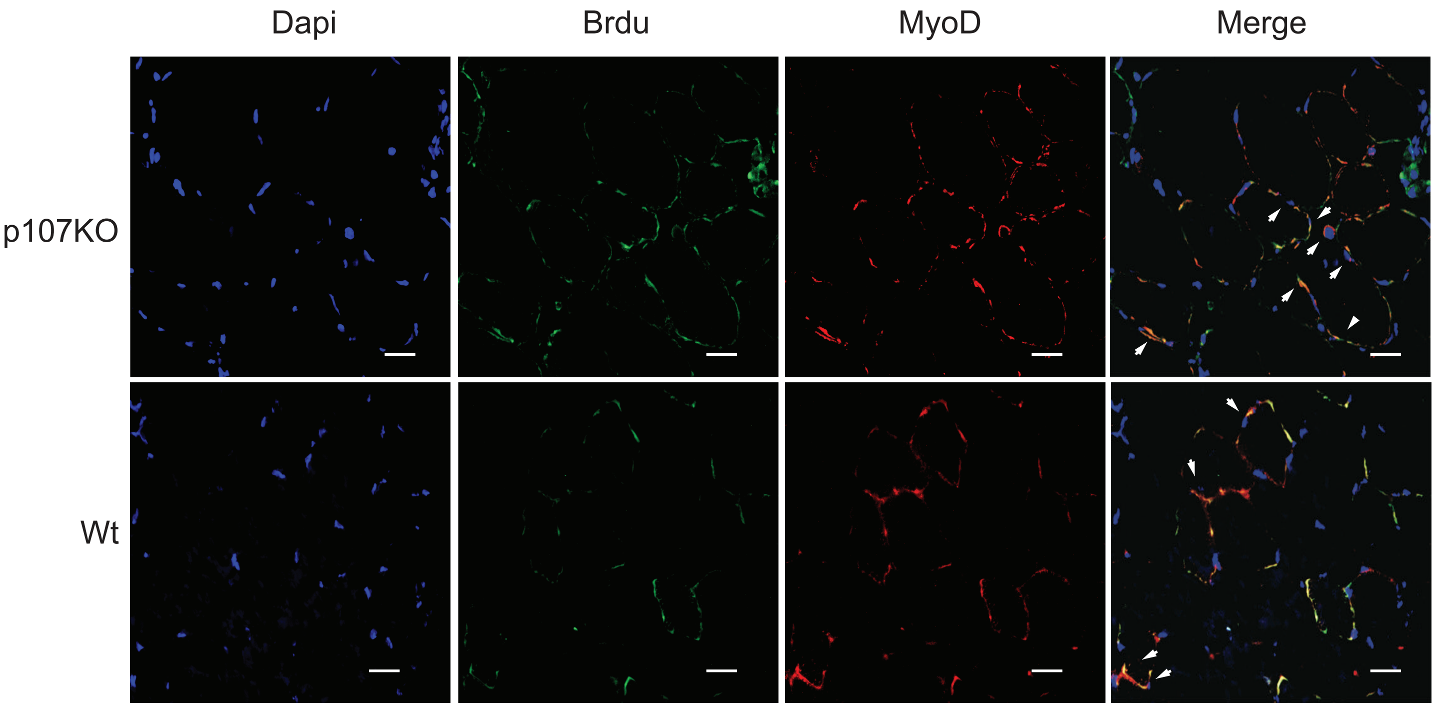


**Suppl. Fig. 11.** Confocal immunofluorescence microscopy for Brdu (green), MyoD (red), Dapi (blue) and Merge of tibialis anterior muscle tissue section from wild type (Wt) and p107 genetically deleted (p107KO) mice 2 days post injection with cardiotoxin that had been treated with bromodeoxyuridine (Brdu) on the previous day (scale bar 20µm).

### Supplemental Figure 12

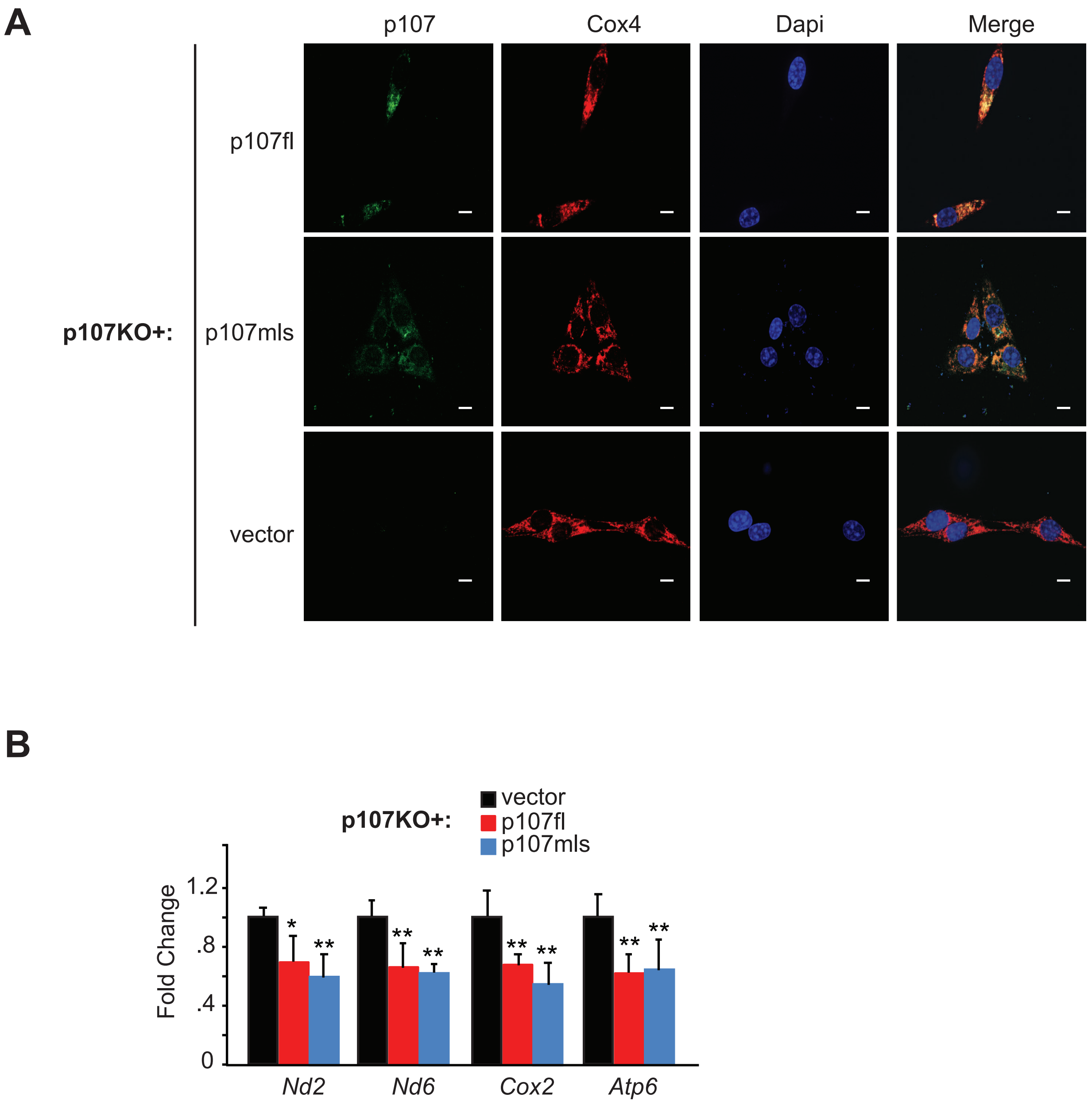


**Suppl. Fig. 12. p107 mitochondrial localization in p107KO MPs down regulates mtDNA gene expression.** p107 genetically deleted (p107KO) c2MPs transfected with either empty vector alone or together with full length p107 (p107fl) or mitochondria localized p107 (p107mls) were assessed for **(A)** confocal immunofluorescence for p107 (green), Cox4 (red), Dapi (blue) and Merge (scale bar 10µm) and **(B)** Gene expression analysis by qPCR of mitochondrial encoded genes *Nd2*, *Nd6*, *Cox2* and *Atp6*, n=4; asterisks denote significance, $*p<0.05$; $**p<0.01$; one way-Anova and post hoc Tukey.

#### Supplemental Figure 13

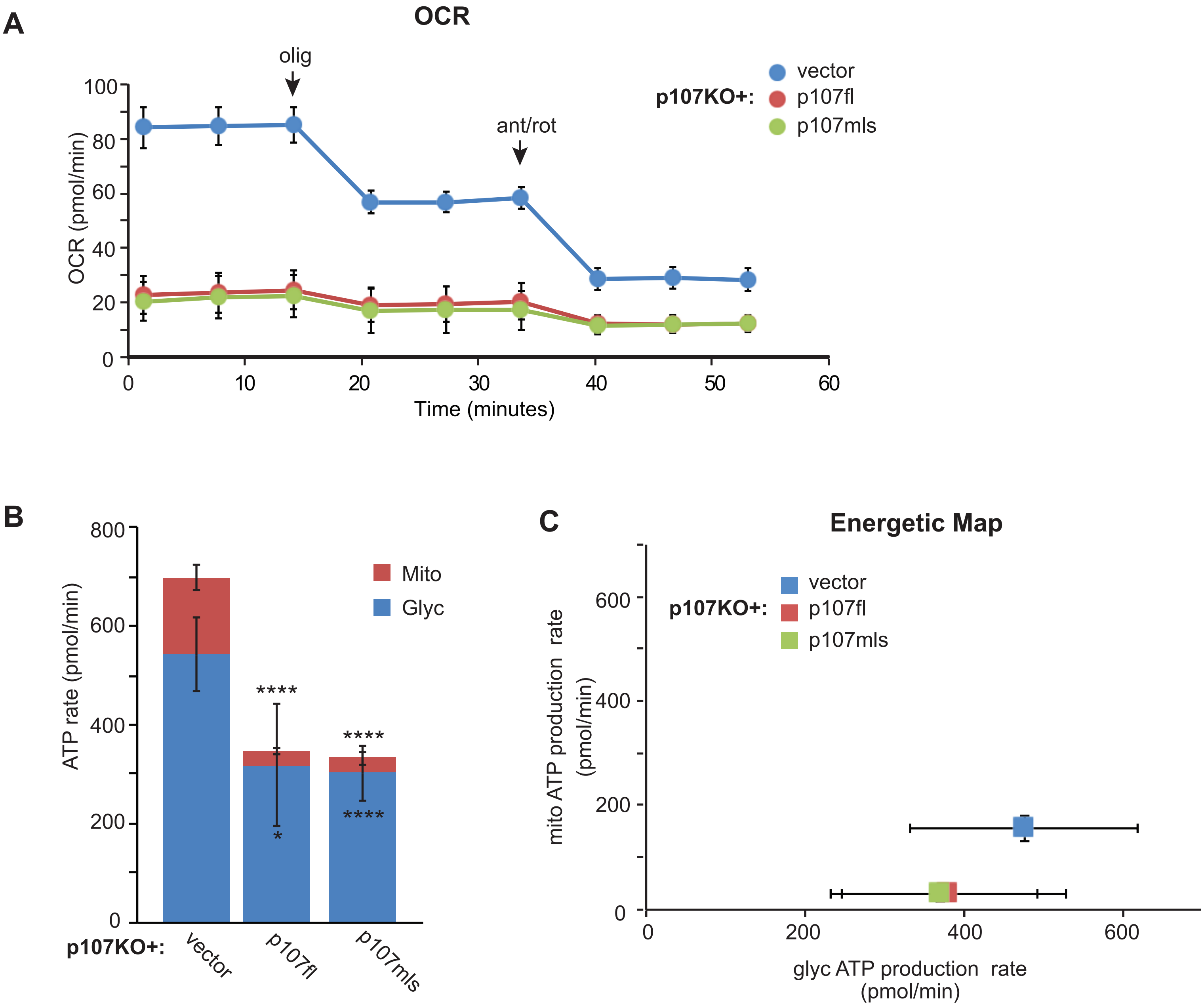


**Suppl. Fig. 13. p107 mitochondrial localization in p107KO MPs reduce production of total and mitochondrial ATP.** Live cell metabolic analysis by Seahorse of **(A)** Oxygen consumption rate (OCR) with addition of oligomycin (olig) and antimycin A (ant), **(B)** ATP production rate from mitochondria (Mito) and glycolysis (Glyc) and **(C)** energetic map for proliferating p107 genetically deleted (p107KO) c2MPs transfected with either empty vector or together with full length p107 (p107fl) or mitochondria localized p107 (p107mls), n=6-8; asterisks denote significance, *$p$<0.05, **$p$<0.01; two-way Anova and post hoc Tukey.

### Supplemental Figure 14

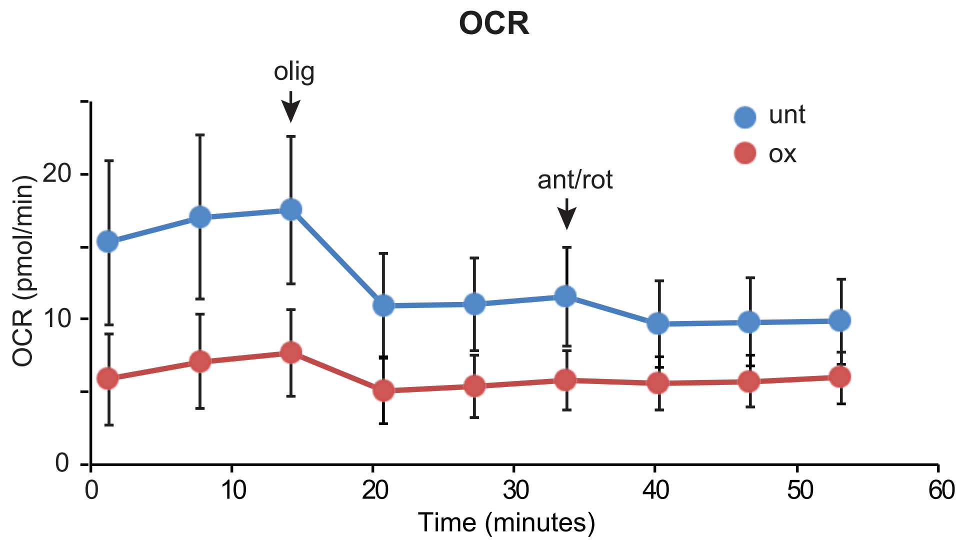


**Suppl. Fig. 14. Oxamate treated cells have reduced production of oxygen consumption rate (OCR).** Live cell metabolic analysis by Seahorse of Oxygen consumption rate (OCR) with addition of oligomycin (olig) and antimycin A (ant) for c2MPs untreated (unt) or treated oxamate (ox), n=6-8.

#### Supplemental Figure 15

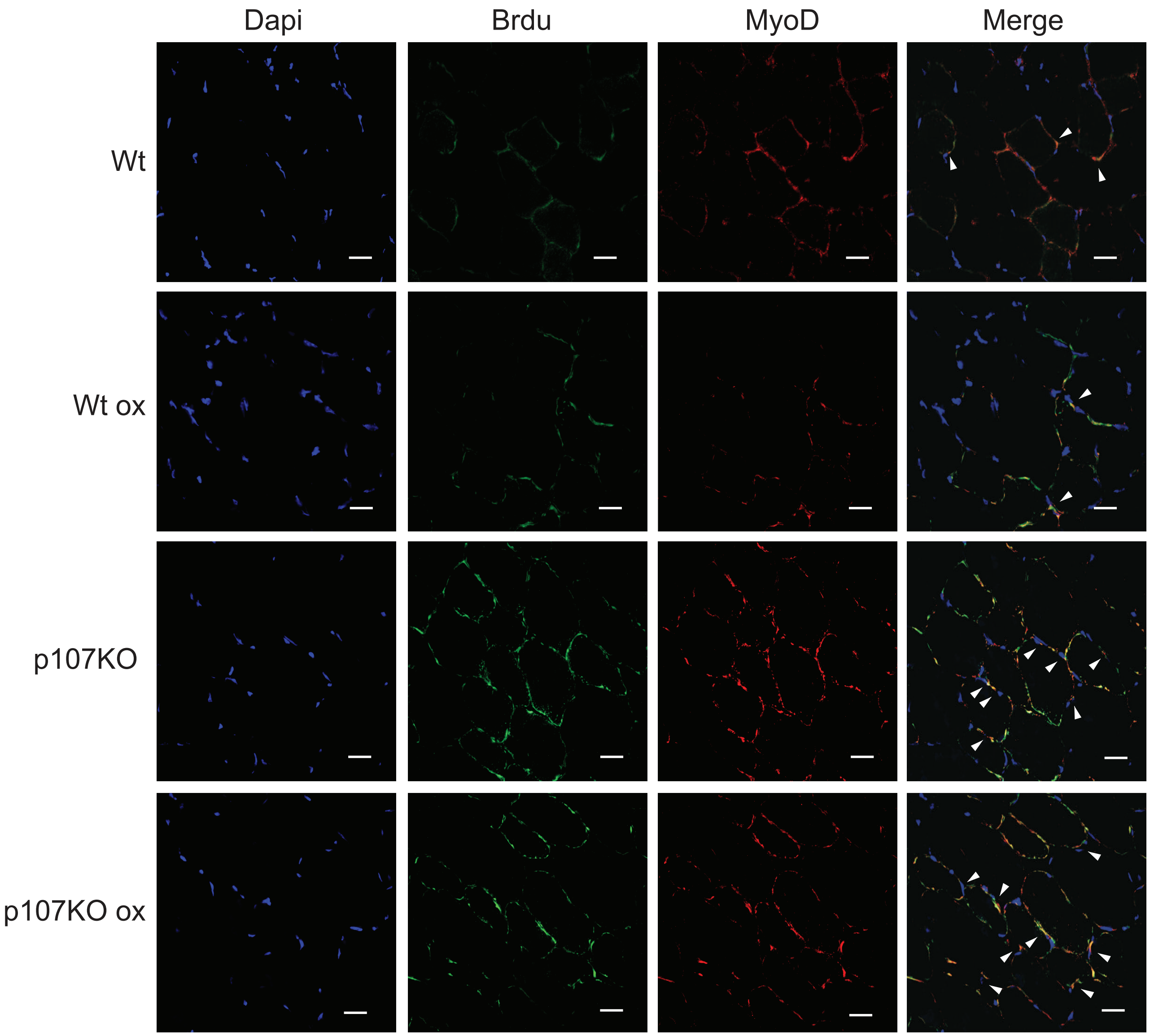


**Suppl. Fig. 15.** Confocal immunofluorescence microscopy for Brdu (green), MyoD (red), Dapi (blue) and Merge of tibialis anterior muscle tissue section from wild type (Wt) and p107 genetically deleted (p107KO) mice 2 days post injection with cardiotoxin that had been treated with bromodeoxyuridine (Brdu) on the previous day in the absence or presence of oxamate (ox) for 4 days (scale bar 20μm).

#### Supplemental Figure 16

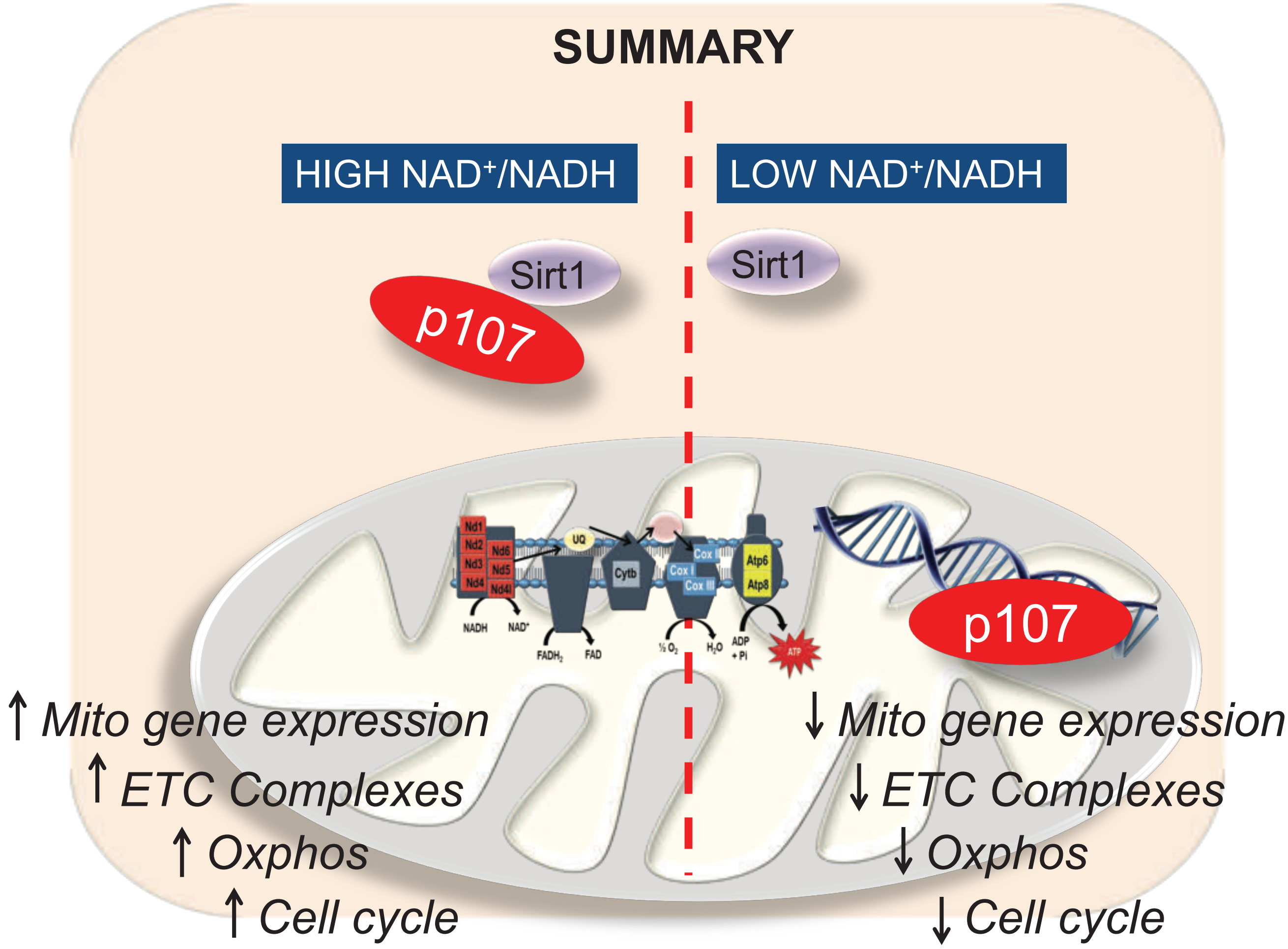


**Suppl Fig. 16. The mitochondrial function of the transcriptional co-repressor, p107 governs muscle progenitor (MP) proliferation based on Sirt1 activity.** When the cytoplasmic $NAD^+$/NADH ratio is high, Sirt1 is active interacting with p107 preventing its mitochondrial localization. This causes de-repression of the mitochondrial encoded genes, thus enhancing Oxphos generation that increases MP proliferation rate. Contrarily, with low $NAD^+$/NADH ratio, Sirt1 is inactive and p107 is free to relocate to the mitochondria, where it binds to the mitochondrial DNA D loop promoter and represses its gene expression. This causes down regulation of mitochondrial Oxphos that decelerates MP cell cycle progression and proliferation rate.

**Supplemental Table 1. Statistical significance values at each timepoint for various ATP generation capacity graphs.**

| A) ATP generation capacity for proliferation (G) and growth arrest (Ga), n=4 (Fig. 2E) | |
|---|---|
| **Time (min.)** | **Significance** |
| 1 | 0.033131343 (*) |
| 2 | 0.016586179 (*) |
| 3 | 0.011167771 (*) |
| 4 | 0.00665779 (**) |
| 5 | 0.005020464 (**) |
| 6 | 0.005420062 (**) |
| 7 | 0.001023739 (***) |
| 8 | 0.002013576 (**) |
| 9 | 0.001840361 (**) |
| 10 | 0.001772278 (**) |
| 11 | 0.001374734 (**) |
| 12 | 0.001369558 (**) |
| 13 | 0.001820507 (**) |
| 14 | 0.000942071 (***) |
| 15 | 0.000710462 (***) |

| B) ATP generation capacity for proliferating control (Ctl) and p107KO c2MPs, n=4 (Fig. 2F) | |
|---|---|
| **Time (min.)** | **Significance** |
| 1 | 0.009417287 (**) |
| 2 | 0.008062857 (**) |
| 3 | 0.005542089 (**) |
| 4 | 0.00526662 (**) |
| 5 | 0.004568636 (**) |
| 6 | 0.008806086 (**) |
| 7 | 0.015920646 (*) |
| 8 | 0.04541628 (*) |
| 9 | 0.018155165 (*) |
| 10 | 0.020427915 (*) |
| 11 | 0.0183923 (*) |
| 12 | 0.020811991 (*) |
| 13 | 0.021514531 (*) |
| 14 | 0.021540274 (*) |
| 15 | 0.003472532 (**) |

| C) ATP generation capacity for c2MPs grown in SM containing 5.5mM and 25mM glucose, n=4, (Fig. 3F) | |
|---|---|
| **Time (min.)** | **Significance** |
| 1 | 0.035331802 (*) |
| 2 | 0.030200259 (*) |
| 3 | 0.025596033 (*) |
| 4 | 0.024064309 (*) |
| 5 | 0.02435088 (*) |
| 6 | 0.022419532 (*) |
| 7 | 0.021898214 (*) |
| 8 | 0.020053804 (*) |
| 9 | 0.017606462 (*) |
| 10 | 0.009207275 (**) |
| 11 | 0.011190924 (*) |
| 12 | 0.036293605 (*) |
| 13 | 0.016513239 (*) |
| 14 | 0.011994536 (*) |
| 15 | 0.005802656 9 (**) |

| D) ATP generation capacity for Ctl and Sirt1KO treated with 5.5mM glucose, n=4 (Fig. 4D) | |
|---|---|
| **Time (min.)** | **Significance** |
| 1 | 0.001807382 (**) |
| 2 | 0.000423724 (***) |
| 3 | 0.004639819 (**) |
| 4 | 0.000760897 (***) |
| 5 | 0.001120419 (**) |
| 6 | 0.000853747 (***) |
| 7 | 0.000975817 (***) |
| 8 | 0.001268889 (**) |
| 9 | 0.000954513 (***) |
| 10 | 0.001572121 (**) |
| 11 | 0.001643781 (**) |
| 12 | 0.001953617 (**) |
| 13 | 0.001685461 (**) |
| 14 | 0.002527076 (**) |
| 15 | 0.002392154 (**) |

| E) ATP generation capacity for c2MPs treated or untreated with 10mM nam, n=4 (Fig. 4H) | |
|---|---|
| **Time (min.)** | **Significance** |
| 1 | 0.000360044 (***) |
| 2 | 0.006963349 (**) |
| 3 | 0.014770086 (*) |
| 4 | 0.02132306 (*) |
| 5 | 0.022632029 (*) |
| 6 | 0.032207431 (*) |
| 7 | 0.034483084 (*) |
| 8 | 0.035670347 (*) |
| 9 | 0.007821973 (**) |
| 10 | 0.020493916 (*) |
| 11 | 0.027989954 (*) |
| 12 | 0.032556922 (*) |
| 13 | 0.038031967 (*) |
| 14 | 0.034633286 (*) |
| 15 | 0.032465285 (*) |

| F) ATP generation capacity for c2MPs treated or untreated with 10μM res, n=4 (Fig. 4N) | |
|---|---|
| **Time (min.)** | **Significance** |
| 1 | 0.064444463 (*) |
| 2 | 0.027493958 (*) |
| 3 | 0.021748346 (*) |
| 4 | 0.016031546 (*) |
| 5 | 0.01277941 (*) |
| 6 | 0.010063891 (**) |
| 7 | 0.006263203 (**) |
| 8 | 0.0040332 (**) |
| 9 | 0.002687763 (**) |
| 10 | 0.00280281 (**) |
| 11 | 0.00231082 (**) |
| 12 | 0.005146623 (**) |
| 13 | 0.005146623 (**) |
| 14 | 0.00451611 (**) |
| 15 | 0.005339423 (**) |

| G) ATP generation capacity for c2MPs treated or untreated with 25µM res, n=4 (Suppl. Fig. 7C) | |
|---|---|
| **Time (min.)** | **Significance** |
| 1 | 0.000959268 (***) |
| 2 | 0.001423853 (**) |
| 3 | 0.00096012 (***) |
| 4 | 0.000862286 (***) |
| 5 | 0.001400103 (**) |
| 6 | 0.001343943 (**) |
| 7 | 0.001396845 (**) |
| 8 | 0.0025641 (**) |
| 9 | 0.003482068 (**) |
| 10 | 0.006695159 (**) |
| 11 | 0.008173165 (**) |
| 12 | 0.010742694 (*) |
| 13 | 0.010742694 (**) |
| 14 | 0.017190247 (*) |
| 15 | 0.019641938 (*) |

**Supplemental Table 2. Primer Sets Used**

| Gene Name | Sequence Accession Number | Amplicon Length (bp) | Forward primer sequence | Reverse primer sequence |
|---|---|---|---|---|
| Rplp0 (36B4) | MGI:1927636 | 29 | GAGGAATCAGATGAGG ATATGGGA | AAGCAGGCTGACTTGG TTGC |
| mt-Nd2 (Nd2) | MGI:102500 | 121 | CATAGGGGCATGAGGA GGACT | TGAGTAGAGTGAGGGA TGGGTTG |
| mt-Nd6 (Nd6) | MGI:102495 | 44 | TGTTGCAGTTATGTTGG AAGGAG | CAAAGATCACCCAGCT ACTACC |
| mt-Co2 (Cox2) | MGI:102503 | 98 | AGTTGATAACCGAGTC GTTCTG | CTGTTGCTTGATTTAGT CGGC |
| mt-Atp6 (Atp6) | MGI:99927 | 55 | TCCCAATCGTTGTAGC CATC | TGTTGGAAAGAATGGA GTCGG |
| D-loop | MF 133498.1 | 173 | GCGTTATCGCCTCATA CGTT | GGTGCGTCTAGACTGT GTG |
| Nfe2l2 (Nrf2) | MGI:108420 | 146 | AGAGCAACTCCAGAAG GAACAG | TGTGGGCAACCTGGGA GTAG |
| Mfn2 | MGI:2442230 | 103 | AGAGGCAGTTTGAGGA GTGC | ATGATGAGACGAACGG CCTC |
| Ppargc1a (Pgc-1α) | MGI:1342774 | 126 | TACGCAGGTCGAACGA AACT | ACTTGCTCTTGGTGGA AGCA |
| Nrf1 | MGI:1332235 | 154 | GTTGGTACAGGGGCAA CAGT | TCGTCTGGATGGTCATT TCA |
| H19 | MGI:95891 | 207 | GTACCCACCTGTCGTC C | GTCCACGAGACCAATG ACTG |
| Mt-Co1 | MGI:102504 | 342 | CCCAATCTCTACCAGC ATC | GGCTCATAGTATAGCT GGAG |
| Slc25a4 (ANT 1) | MGI:1353495 | 174 | GTCTCTGTCCAGGGCA TCAT | ACGACGAACAGTGTCA AACG |
