## Supplemental Materials and Methods for "p107 mediated mitochondrial function controls muscle stem cell proliferative fates"

The c2c12 myogenic progenitor cell line (c2MP) was purchased from the American Tissue Type Culture (ATTC) and grown in Dulbecco's Modified Eagle Medium (DMEM) (Wisent) containing 25mM glucose supplemented with 10% fetal bovine serum (FBS) and 1% penicillin streptomycin. Primary myogenic progenitor cells (prMPs) were grown on rat tail collagen I (ThermoFisher Scientific) coated dishes containing in Ham's F10 Nutrient Mix Media (ThermoFisher Scientific) supplemented with 20% FBS, 1% penicillin streptomycin and 2.5ng/ml bFGF (PeproTech).

For the nutrient specific experiments, c2MPs or prMPs were grown in stripped DMEM with 10% FBS and 1% penicillin streptomycin for 20 hours supplemented with 1mM, 5.5mM or 25mM glucose; or 4mM or 20mM glutamine; or 4mM glutamine with 25mM glucose or 10mM galactose for 6 hours. For drug specific treatment, c2MPs were grown in DMEM with 10% FBS, 1% penicillin streptomycin, supplemented with 2.5mM oxamate (ox) for 40 hours. For Sirt1 inhibition, cells were grown in stripped DMEM with 10% FBS and 1% penicillin streptomycin and 5.5mM glucose with or without 10mM nicotinamide (nam) for 20 hours. For Sirt1 activation, cells were grown in DMEM with 10% FBS, 1% penicillin streptomycin with or without 10µM resveratrol (res) and for inactivation with 25µM res for 18 hours.

**Primary myogenic progenitor cell (prMP) isolation**

All animal experiments were performed following the guidelines approved by the Animal Care Committee of York University. For derivation of prMPs and tissue immunofluorescence wild type and p107KO mice were from M. Rudnicki(LeCouter et al. 1998) maintained on a mixed (NMRI, C57/Bl6, FVB/N) background(Chen et al. 2004). Extensor digitorum longus muscles of 3-month aged mice were dissected from tendon to tendon and digested in filter sterilized 0.2% type 1 collagenase (Sigma Aldrich) in serum free DMEM for 30 minutes. Upon unravelling, it was transferred to a pre-warmed petri dish containing DMEM (Wisent) with 1% penicillin, streptomycin. The muscle was then flushed gently with the media until it released the fibers with an intermittent incubation at 37°C every 5 minutes. After 30 minutes, individual fibers were transferred into 24 well rat tail collagen I (ThermoFisher Scientific) coated tissue culture plate containing pre-warmed Ham's F10 Nutrient Mix Media (ThermoFisher Scientific) with 20% FBS, 1% penicillin, streptomycin and 2.5ng/ml bFGF (PeproTech). After 3 days, the fibers were transferred to collagen coated Ham's F10 Nutrient Mix Media with 20% FBS, 1% penicillin, streptomycin and 2.5ng/ml bFGF (PeproTech). The media was changed every alternate day, until the fibers generated prMPs that were collected by trypsinization and passaging onto collagen coated plates to remove live fibers.

**Cloning**

The p107mls expression plasmid that expresses p107 only in the mitochondria was made by cloning full length p107 into the pCMV6-OCT-HA-eGFP expression plasmid vector(Newman et al. 2016) that contains a mitochondrial localization signal. We used the following forward 5'-CACCAATTGATGTTCGAGGACAAGCCCCAC-3' and reverse 5'-CACAAGCTTTTAATGATTTGCTCTTTCACT-3' primer sets that contain the restriction sites Mfe1 and HindIII, respectively, to amplify full a length p107 insert from a p107 Ha tagged plasmid(Iwahori et al. 2017). The restriction enzyme digested full length p107 insert was then ligated to an EcoRI/HindIII digest of pCMV6-OCT-HA-eGFP, which removed the HA-eGFP sequences but retained the n-terminal mitochondrial localization signal (OCT).

**Transfections**

The calcium chloride method was used for transfections, whereby a mixture containing a total of 10μg of plasmid DNA, 125mM $CaCl_2$ and $H_2O$ were added dropwise to HEBS buffer (274mM

NaCl, 10mM KCl, 1.4mM $Na_2HPO_4$, 15mM D-glucose and 42mM HEPES), incubated at room temperature for 1 hour and then added to $1 \times 10^5$ cells that had been passaged on the previous day. 18 hours post transfection, fresh DMEM containing 25mM glucose supplemented with 10% fetal bovine serum (FBS) and 1% penicillin streptomycin was added, and the cells used the next day. For overexpression studies, at least 4 different p107KO c2MPs were transfected as above with GFP mitochondrial localization empty vector pCMV6-OCT-HA-eGFP(Newman et al. 2016), p107fl expressing full length p107 tagged HA, and p107mls expressing full length p107, which is directed to the mitochondria. For Sirt1 overexpression experiments, c2MPs were transfected with p107fl alone expressing full length p107 tagged HA or with full length (Sirt1fl) or dominant negative (Sirt1dn) Sirt1(Boily et al. 2009).

**p107KO and SirtKO cell line derivation**

Crispr/Cas9 was used to generate p107 and Sirt1 genetically deleted c2MP (p107KO and SirtKO) cell lines. For p107KO c2MPs, c2c12 cells were simultaneously transfected with 3 pLenti-U6-sgRNA-SFFV-Cas9-2A-Puro plasmids each containing a different sgRNA to target p107 sequences 110 CGTGAAGTCATCCAGGGCTT, 156 GGGAGAAGTTATACACTGGC and 350 AGTTTCGTGAGCGGATAGAA (Applied Biological Materials), and for Sirt1KO with 2 Double Nickase plasmids each containing a different sgRNA to target sequences 148 CGGACGAGCCGCTCCGCAAG and 110 CCATGGCGGCCGCCGCGGAA (Santa Cruz Biotechnology). Following 18 hours of incubation, the media was changed for 6 hours and then passaged into 96 well tissue culture plates. On the next day, the media was aspirated and replenished with DMEM containing 2mg/ml puromycin for antibiotic selection. Following, the media was changed every 2 days until cell clones were visible. Any surviving clones was grown and tested by Western blotting. For control cells, c2MPs cells were transfected by empty pLenti-U6-sgRNA-SFFV-Cas9-2A-Puro (Applied Biological Materials) and selected as above.

**Mitochondrial, nuclear and cytosolic isolation**

For nuclear and cytoplasmic isolation, at least cells were pelleted, washed in PBS, dissolved in 500μl of cytoplasmic buffer (10mM Tris pH 7.4, 10mM NaCl, 3mM $MgCl_2$, 0.5% NP-40 and protease inhibitors) and incubated on ice for 5 minutes followed by rocking on ice for 5 minutes. After centrifugation at 2500g for 5 minutes at 4ºC, the supernatant was stored as the "cytoplasmic fraction". The cell pellet that represents the "nuclear fraction" was then washed 8 times with the cytoplasmic buffer and lysed with nuclear lysis buffer containing 50mM Tris pH 7.4, 5mM $MgCl_2$, 0.1mM EDTA, 1mM dithiothreitol (DTT), 40% (wt/vol) glycerol and 0.15 unit/μl benzonase (Novagen).

For mitochondrial and cytosolic isolation, at least 1 million cells were washed in PBS, pelleted, dissolved in 5 times the packed volume with isolation buffer (0.25M Sucrose, 0.1% BSA, 0.2mM EDTA, 10mM HEPES, pH 7.4; with 1mg/ml of each pepstatin, leupeptin and aprotinin protease inhibitors), transferred into a prechilled Dounce homogenizer and homogenized loose (5-6 times) and tight (5-6 times) on ice. The homogenate was transferred into an eppendorf tube and centrifuged at 1000g at 4ºC for 10 minutes. The supernatant was then centrifuged at 14000g for 15 min at 4ºC and the resulting supernatant was saved as "cytosolic fraction". The pellet representing the "mitochondrial fraction" was washed twice and dissolved in 50μl of isolation buffer. The mitochondria were lysed by repeated freeze-thaw cycles (3 times each) on dry ice.

**Mitochondria fractionation**

Mitochondrial fractions were isolated using a hypotonic osmotic shock approach(Lu et al. 2009). For this, cells were collected by centrifugation at 1500rpm and the cell pellet resuspended in STE buffer (250mM sucrose, 5mM Tris pH 7.4 and 1mM EGTA), Dounce homogenized and centrifuged at 1000g for 3 mins to remove cell debris. The supernatant was then centrifuged at 10,000g for 10 mins to isolate pelleted mitochondria. The mitochondria were resuspended in STE buffer containing 250mM sucrose and centrifuged at 16000g for 10 minutes to isolate matrix (M) and inner membrane (IM) proteins and the outer membrane (OM) proteins in the pellet and supernatant, respectively. To obtain a pure mitochondria matrix protein (M) fraction the isolated mitochondria were resuspended in hypotonic STE buffer containing 25mM sucrose and centrifuged at 16000g for 10 mins. After centrifugation, the soluble inner membrane (IMS) and outer membrane (OM) protein fractions were present in the supernatant and the matrix protein fraction (M) in the pellet. To demonstrate mitochondria matrix veracity, 50μg/ml porcine trypsin (Promega) was added for 30 mins followed by 20μg/ml trypsin inhibitor aprotinin (Roche) for 10 minutes before centrifugation at 16000g for 10 mins. Only fractions containing M proteins were unavailable for trypsin digestion. As a control whole cell treatment of c2MPs with both buffers showed that trypsin treatment can digest p107.

**Western blot analysis**

For Western blot analysis, cells were lysed in RIPA buffer (0.5% NP-40, 0.1% sodium deoxycholate, 150mM NaCl, 50mM Tris-Cl pH 7.5, 5mM EDTA) or mitochondrial isolation buffer (0.25M sucrose, 0.1% BSA, 0.2mM EDTA, 10mM HEPES, pH 7.4) containing protease inhibitors (1mg/ml of each of pepstatin, aprotinin and leupeptin). Protein lysates were loaded on gradient gels (6-15%) or 7.5% gels. Proteins were transferred using a wet transfer method onto a 0.45μm pore sized polyvinylidene difluoride membrane (Santa Cruz Biotechnology) at 4ºC for 80 minutes at 100V. The membranes were blocked for an hour at room temperature in 5% non-fat milk in Tris-Buffered saline (TBS-150mM NaCl and 50mM Tris base) containing 0.1% Tween-20 (TBST). The membranes were probed overnight at 4ºC with primary antibodies (listed below) diluted in 5% non-fat milk or 1% BSA in TBST. The membranes were then washed three times with TBST and secondary antibodies conjugated with horseradish peroxidase diluted in 5% non-fat milk in TBST were added for an hour at room temperature with gentle rocking. The membranes were then washed 3 times with TBST and visualized with chemiluminescence on photographic films. Protein levels were evaluated by densitometry using Image J software.

**Primary antibodies used**

α-tubulin (66031, Proteintech); Cox4 (ab16056, Abcam) and Total OXPHOS rodent WB antibody cocktail (Abcam); p107-C18, p107-SD9, histone H3-C16, Sirt1-B7, IgG-D7, Ha-tag-F7, Brdu-MoBU-1, Pax-7 (EE8) (Santa Cruz Biotech); MyoD (Novus Biologicals) and Sirt1-D1D7 (Cell Signaling).

**Co-immunoprecipitation**

For Immunoprecipitation (IP), protein lysates were pre-cleared with 50µl protein A/G plus agarose beads (Santa Cruz Biotechnology) by rocking at 4ºC for an hour. The sample was centrifuged at 15000 rpm for a minute. Fresh protein A/G agarose beads along with 5µg of p107-C18, Sirt1-B7 or IgG-D7 antibody (Santa Cruz Biotechnology) antibody were added to the supernatant and rocked overnight at 4ºC. The next day the pellets were washed 3 times with wash buffer (50mM HEPES pH 7.0, 250mM NaCl and 0.1% Np-40) and loaded onto polyacrylamide gels and Western blotted for p107-SD9 (Santa Cruz Biotechnology) or Sirt1-D1D7 (Cell Signaling). Inputs represent 10% of lysates that were immunoprecipitated.

**qPCR**

RNA was isolated using Qiazol reagent (Qiagen) and concentration was determined by NanoDrop 2000 (ThermoFisher Scientific). qPCR experiments were performed according to the MIQE (Minimum Information for Publication of Quantitative Real-Time PCR Experiments) guidelines (Bustin et al. 2009). The optical density (OD) of RNA was measured using the NanoDrop 2000 (Thermo Fisher Scientific), RNA purity was inferred by the A260/280 ratio (~1.80 is pure). 1μg of RNA was reverse transcribed into cDNA using the All-in-One cDNA Synthesis SuperMix (Bimake) and the cDNA used for qPCR. qPCR assays were performed on Light cycler 96 (Roche) using SYBR green Fast qPCR Master mix (Bimake) with appropriate primer sets and Rplp0 (36B4) as a normalization control was used (**Suppl. Table 2**). Relative expression of cDNAs was determined with 36B4 as the internal control using the ΔΔCt method. For fold change, the ΔΔCt was normalized to the control. Student t-tests, one-way or two-way Anova and Tukey post hoc tests were used for comparison and to obtain statistics.

**Mitochondrial and nuclear DNA content**

To obtain the mitochondrial to nuclear DNA ratio (mtDNA/nDNA), cells grown on a 6cm tissue culture plate were untreated or treated with 1mM 5-aminoimidazole-4-carboxamide-1-β-D-ribofuranoside (aicar) (Toronto Research Chemicals) in presence of 5.5mM or 25mM glucose, with or without 10mM Nam or 10μM resveratrol for 24 hours. The cells were washed in PBS and lysed with 600μl of lysis buffer containing 100mM NaCl, 10mM EDTA, 0.5% SDS solution, 20mM Tris HCl; pH 7.4 and 6μl of 1μg/μl proteinase k (ThermoFisher Scientific). Following incubation at 55ºC for 3 hours, 100μg/ml RNase A (ThermoFisher Scientific) was added at 37ºC for 30 minutes. After, 250μl of 7.5M ammonium acetate and 600μl of isopropanol were added, the cells were centrifuged at 15000g for 10 minutes at 4ºC. The supernatant was discarded and the pellet containing mitochondrial and nuclear DNA was washed with 70% ethanol and resuspended in Tris EDTA buffer (10mM Tris HCl pH 7.4, 1mM EDTA). qPCR assays were performed on 10ng DNA with primer sets to Mt-Co1 and H19 (**Suppl. Table 2**) representing total mitochondrial and nuclear DNA content, respectively. Ct values were obtained and the ratio mtDNA:nDNA were determined by the formula: nDNA Ct- mtDNA Ct.

**Quantitative chromatin immunoprecipitation assay (qChIP)**

qChIP was performed on mitochondrial lysates containing only mitochondrial DNA. Mitochondrial fractions were collected as described above, washed twice in PBS by centrifugation at 14000g for 15 minutes at 4°C, resuspended in 200µl of PBS containing 1% formaldehyde and rocked at room temperature for 30 minutes to fix the cells. The fixation reaction was quenched by adding 125mM of glycine in PBS and rocked for 5 minutes at room temperature. The fixed pellet was washed twice in PBS containing 100mM NaF and 1mM $Na_3VO_4$ by centrifuging at 14000g for 5 minutes at 4°C. The pellet was the resuspended in 500µl of ChIP lysis buffer (40mM Tris, pH 8.0, 1% Triton X-100, 4mM EDTA, 300mM NaCl) and sonicated at 24% amplitude, 15 seconds on, 15 second off for 3 cycles (Model 120 Sonic Dismembrator, ThermoFisher Scientific). Following sonication, the samples were centrifuged at 13000 rpm for 10 minutes at 4°C and the supernatant was transferred to a new tube containing 100µl of dilution buffer 1 (40mM Tris, pH 8.0, 4mM EDTA) from which the input controls were removed before 150 µl of dilution buffer 2 (40mM Tris, pH 8.0, 0.5% Triton X-100, 4mM EDTA, 150mM NaCl) was added. To preclear, 50µl of protein A/G agarose beads (Santa Cruz Biotechnology) was added and rocked for 90 minutes at 4°C. The beads were then pelleted and discarded, and to the supernatant was added 5µg of p107 antibody (p107- C-18) (Santa Cruz Biotechnology) or IgG antibody (IgG-D-7) (Santa

Cruz Biotechnology). This was rocked overnight at 4°C and on the following day, 50μl of protein A/G agarose beads were added and rocked for 90 minutes at 4°C. The beads were collected by centrifugation and were washed sequentially with 5 minutes rocking at 4°C before centrifugation by adding the following: low salt complex wash buffer (0.1% SDS, 1% Triton X-100, 2mM EDTA, 20mM Tris HCl, pH 8.0, 150mM NaCl), high salt complex wash buffer (0.1% SDS, 1% Triton X-100, 2mM EDTA, 20mM Tris HCl, pH 8.0, 500mM NaCl), LiCl wash buffer (0.25M LiCl, 1% NP-40, 1% deoxycholic acid, 1mM EDTA, 10mM Tris, pH 8.0) and 2 washes with TE buffer (10mM Tris HCl, pH 8.0, 1mM EDTA). After the last wash, the mtDNA-protein complexes were isolated by resuspending the beads in 250μl of elution buffer (1% SDS, 0.1M $NaHCO_3$), vortexing, rocking for 15 minutes at room temperature and centrifuging at 2000 rpm in a microcentrifuge for 2 minutes. The supernatant was transferred to a clean tube and to the remaining beads another 250μl of elution buffer was added and the isolation step repeated. For isolation of mtDNA fragments, to the 500μl of mtDNA-protein complexes in elution buffer, 20μl of 5M NaCl was added and incubated at 65°C overnight. The next day, the DNA was isolated using DNA purification kit (Thermo Fisher Scientific), and the concentration determined using the NanoDrop 2000 (ThermoFisher Scientific). Relative occupancy was determined by amplifying isolated DNA fragments using the D-loop primer sets (**Suppl. Table 2**) analyzed using the ΔΔCt method and the fold changes were normalized to IgG ΔΔCt values.

**$NAD^+$/NADH assay**

Cells growing in 6cm tissue culture plates were washed with PBS and scraped into 500μl of 0.5M perchloric acid, vortexed and freeze-thawed on dry ice three times. The cells were then centrifuged at 4°C for 5 minutes at 7,000 rpm in a microfuge and 100μl of 2.2M $KHCO_3$ was added to the supernatant on ice. This was again centrifuged at 7,000 rpm for 15 minutes at 4°C and the supernatant was collected to be analyzed. For the lactate assay; 20µl of the supernatant, 258µl of lactate buffer (1M glycine, 500mM hydrazine sulfate, 5mM EDTA), 20µl of 25mM β-nicotinamide-adenine dinucleotide (NAD) (Roche) and 2µl of porcine heart lactate dehydrogenase (Ldh) (Sigma-Aldrich) were added to each well of an assay plate. For the pyruvate assay; 20µl of the supernatant, 218µl of pyruvate buffer (1.5M Tris, pH 8), 180µl of 6µM β-nicotinamide-adenine dinucleotide, reduced (NADH) (Roche) and 2µl of rabbit skeletal muscle LDH (Sigma-Aldrich) were added for each well. The assay plates were then read by a microplate reader (Glomax, Promega) with excitation at 340nm and emission peak at 450nm and the concentration values were attained by comparing to standard curves. The standard curves were made with NADH (Roche) at 0, 2, 3, 4, 5, 6, 7, and 10mM in lactate buffer for the lactate assay and in pyruvate buffer for the pyruvate assay. Estimation of free $NAD^+$/NADH in cells was based on lactate/pyruvate (pyruvate + NADH + $H^+$ = lactate + $NAD^+$). The results were normalized to control samples and graphed.

**ATP generation capacity and rate assay**

The ATP generation capacity assay is based on the requirement of luciferase for ATP in producing light from the reaction:

$$\text{Luciferin} + \text{ATP} + O_2 \xrightarrow{\text{Luciferase}} \text{Oxyluciferin} + \text{AMP} + \text{Pyrophosphate} + CO_2 + \text{Light}$$

It was measured from mitochondrial fractions by using the ATP determination kit (ThermoFisher Scientific). In each well of a 96 well assay plate, 10µl of the isolated mitochondria was added to 4µl of 100mM ADP and 86 µl of reaction mix (1.78µl $ddH_2O$, 100µl of 20X reaction buffer, 20µl of 0.1M DTT, 100µl of 10mM D-luciferin and 0.5µl firefly luciferase from a 5mg/mL stock solution, all supplied by the kit manufacturer). The plate was then read over a period of 15 minutes every 1 minute for emission peak at 560 nm by a microplate reader (Glomax, Promega). The amount of ATP generated was normalized to the protein content of the mitochondrial fractions determined by Bradford Assay Kit (Biobasic) and graphed over time. The ATP generation rate was calculated from the slope of the graph of ATP that was produced over time up to the point where its production reached a steady state.

**Immunocytochemistry and confocal imaging**

For confocal microscopy, cells were grown on Nunc Lab-Tek™ II chambered tissue culture plates (ThermoFisher Scientific), fixed for 5 minutes with 95% methanol and permeabilized for 30 minutes at 4ºC with blocking buffer (3% BSA and 0.1% saponin in PBS). Cells were then incubated with primary antibody p107-SD9 or HA-tag-F7 (Santa Cruz Biotechnology) in 1:100 dilution in blocking buffer for 1 hour. After 3 washes with 0.05% saponin in PBS (SP), cells were incubated with secondary antibody donkey anti-mouse IgG NL493-conjugated (R and D Systems) at a 1:200 dilution in blocking buffer. Cells were then washed 3 times in SP and re-incubated with primary antibody Cox4 (Abcam) at a 1:100 dilution in blocking buffer for 1 hour. After 3 washes in SP, cells were incubated with secondary antibody goat anti rabbit IgG Alexa Fluor 594 (Thermo FisherScientific) in 1:200 dilution for 1 hour. After, washing 3 times with SP, 4′,6-diamidino-2-phenylindole (Dapi) was added and Vectashield mounting media (Vector) was added before placing the coverslip. Confocal images and Z-stacks were obtained using the Axio Observer.Z1 microscope with alpha Plan-Apochromat 63x/Oil DIC (UV) M27 (Zeiss). Digital images were captured using Axiocam MR R3 (Zeiss). Optical sections were then "stacked" or merged to create high resolution "z-series" images. Z-stack images were also portrayed as orthogonally projected on XY, YZ and XZ plane with maximum intensity using ZEN imaging software (Zeiss). A line was drawn through a representative cell to indicate relative intensity of RGB signals.

**Mitochondrial length and area measurement**

Live cell imaging was used to measure mitochondrial length and area of **c**ells grown on 35mm high glass-bottom µ-Dish tissue culture plates (MatTek Corp). The cells were washed with PBS and stained with 1µl of MitoView red (Biotium) in 5ml serum free DMEM for 30 minutes. The cells were then washed with PBS and refed with serum free DMEM and immediately live cell imaged using the Axio Observer.Z1 (Zeiss) microscope with alpha plan-apochromat 40x/Oil DIC (UV) M27 (Zeiss) in an environment chamber (5% $CO_2$; 37°C). Digital images were captured using Axiocam MR R3 (Zeiss). Mitochondrial length was measured by tracing the mitochondria from one end to the other with a line that was calibrated to the scale bar using Image J software. Area was measured using Image J software according to Ouellet et al (Ouellet et al. 2017). Briefly, threshold was used to select the area of each mitochondria. The background was eliminated and using 'analyze particle', the area of each mitochondria was measured by considering the scale bar as the reference point. At least 100 mitochondria were measured for length and area from each of 3 different controls (Ctl) and p107KO c2MP cell lines. The measurements were categorized and graphed within different intervals of lengths and areas, respectively.

**Cardiotoxin-induced muscle regeneration**

Three-month-old anesthetized wild type and p107KO mice were injected intramuscularly in the tibialis anterior (TA) muscle with 40µl of cardiotoxin (ctx) Latoxan (Sigma) that was prepared by dissolving in water to a final concentration of 10µM. A day after ctx injury, bromodeoxyuridine (Brdu) at 100mg/kg was injected intraperitonially. TA muscles were collected on day 2 post ctx injection, immersed in a 1:2 ratio of 30% sucrose:optimal cutting temperature compound (ThermoFisher Scientific) solution and frozen slowly in liquid nitrogen-cooled isopentane. Mice were also untreated or treated with 750mg/kg ox for four consecutive days, with ctx on the third day and brdu on the fourth day, before the TA muscles were dissected on the fifth day for freezing.

**Immunohistochemistry**

The frozen samples were cross sectioned at 10μm thickness on a cryostat and mounted on positive charged slides (FroggaBio). Muscle tissue sections were washed with PBS and fixed with 4% paraformaldehyde (PFA) for 15 minutes at room temperature. 2N HCl was added for 20 minutes followed by 40mM sodium citrate for 10 minutes at room temperature. After washing in PBS, muscle sections were blocked in blocking buffer (5% goat serum, 0.1% Triton X in PBS) for 30 minutes. Muscle tissue sections were then incubated with primary antibody anti-MyoD (Novus Biologicals) in blocking buffer in 1:100 ratio overnight or p107-SD9 or Pax7-EE8 in blocking buffer for 90 minutes. After three washes in 0.05% Tween 20 in PBS (PBST), cells were incubated with secondary antibody goat anti rabbit IgG Alexa Fluor 594 (ThermoFisher Scientific) or donkey anti-mouse IgG NL493-conjugated (R and D Systems) for one hour in 1:200 ratio. After three washes in PBST the tissue section was re-incubated with primary antibody Brdu-MoBU-1 (Santa Cruz Biotechnology) or Cox4 (Abcam) in blocking buffer in 1:100 ratio for 1 hour. After washing 3 times in PBST, the tissue section was incubated with secondary antibody donkey anti-mouse IgG NL493-conjugated (R and D Systems) or goat anti rabbit IgG Alexa Fluor 594 (ThermoFisher Scientific) in 1:200 ratio for 1 hour. After washing 3 times in PBST, Dapi was added and Vectashield mounting media (Vector) was added before placement of coverslip. The sample was imaged using confocal microscopy with the Axio Observer.Z1 microscope with alpha Plan-Apochromat 40x/Oil DIC (UV) M27 (Zeiss). Pax7 positive proliferating MPs containing p107 co-localized with Cox4 were identified in successive tissue sections that were cut at 6μm instead of 10μm. Proliferating MPs were determined by enumerating the MyoD$^{+}$Brdu$^{+}$Dapi$^{+}$ cells as a percentage of all the Dapi$^{+}$ cells per field. Six fields per muscle section from 4 to 5 different mice for each treatment was used for the analysis.

**Growth curve and Proliferation Rate**

3000 cells were plated for both control and p107KO c2MPs that were untreated or treated with 2.5mM ox. On each of the following 3 days, the number of cells were counted and graphed. Proliferation rate was calculated from the slope of the cell proliferation over time.

**Flow Cytometry**

For cell cycle analysis 50000, Ctl and p107KO cells were treated or untreated with 2.5mM ox for 40hrs or Ctl and Sirt1 KO cells were grown in 5.5mM glucose, to activate Sirt1, for 20 hours or p107KO cells 24 hours post transfection with pCMV6-OCT-HA-eGFP alone or together with p107fl or p107mls were used. The cells were washed twice with PBS by centrifuging at 1200g for 5 minutes and re-dissolved in 1ml of PBS. Cells were then fixed by adding the cell suspension dropwise to 9ml of 70% ethanol while vortexing and then kept at -20°C overnight. The next day, the fixed cells were pelleted at 4000 rpm for 5 minutes and resuspended with 3ml PBS on ice for 10 minutes to rehydrate the cells. The rehydrated cells were centrifuged at 4000 rpm for 5 minutes and dissolved in 500µl of PBS containing 50µg/ml propidium iodide (ThermoFisher Scientific) and 25µg/ml RNAse (ThermoFisher Scientific) before loading on the Attune Nxt Flow Cytometer (ThermoFisher Scientific). Forward and side scatter were appropriately adjusted and propidium iodide was excited with the 488-nm laser and detected in the BL2 channel. The excitation was analyzed by the ModFit LT™ software that provided the percentage of cells present in each of G1, S and G2 phases of cell cycle, which were represented graphically. Flow cytometry for cell cycle analysis of transfected cells consisted of cells transfected with empty vector pCMV6-OCT-HA-eGFP expressing GFP alone or along with p107fl expressing full length p107 or with p107mls expressing full length p107, which is directed to the mitochondria. GFP positive cells were first sorted using the BL1 channel of the cytometer by adjusting forward and side scatter, followed by detection of propidium iodide that was excited with the 488-nm laser and detected in the BL2 channel The excitation was analyzed by the ModFit LT™ software that provided the percentage of cells present in each of G1, S and G2 phases of cell cycle and represented graphically.

**Live cell ATP analysis (Seahorse)**

3000 control or p107KO cells or p107KO cells 24 hours post transfection with pCMV6-OCT-HA-eGFP alone or together with p107fl or p107mls were seeded in DMEM (Wisent) containing 10% FBS and 1% penicillin streptomycin on microplates (Agilent Technologies) and treated according to the required experiment. For analysis, cells were washed in XF assay media supplemented with 10mM glucose, 1mM pyruvate and 2mM glutamine (Agilent Technologies) and incubated in a $CO_2$ free incubator in the same media at 37°C for 1 hour. The media was then removed and replenished with fresh XF media supplemented with 10mM glucose, 1mM pyruvate and 2mM glutamine and the microplate was assessed using the Seahorse XF real-time ATP rate assay kit (Agilent Technologies) on a Seahorse XFe96 extracellular flux analyzer (Agilent Technologies), with addition of 1.5µM of oligomycin and 0.5µM rotenone + antimycin A as per the manufacturer's direction. The energy flux data in real time was determined using Wave 2.6 software.

**Statistical analysis**

Statistical analysis was performed by GraphPad Prism. Student's t-tests were used unless otherwise stated. Results were considered to be statistically significant when $p < 0.05$. Specific data was analyzed using an appropriate one-way or two-way analysis of variance (ANOVA) with a criterion of $p < 0.05$. All significant differences for ANOVA testing were evaluated using a Tukey post hoc test. All data are mean ± SD.
